# Ethnic Identity Profiles Among Adolescents in the ABCD Study: Associations with Resting State Functional Connectivity and Perceived Discrimination

**DOI:** 10.1101/2025.08.24.671805

**Authors:** Taylor R. Jancetic, Micaela Lembo, Chloe L. Hampson, Donisha D. Smith, Julio A. Peraza, Erin Thompson, Mariana Sanchez, Raul Gonzalez, Alan Meca, Angela R. Laird

**Affiliations:** Department of Psychology, Florida International University, Miami, FL, USA; FIU Embrace Center for Advancing Inclusive Communities, Florida International University, Miami, FL, USA; Department of Epidemiology, Florida International University, Miami, FL, USA; Department of Physics, Florida International University, Miami, FL, USA; Center for Children and Families, Florida International University, Miami, FL, USA; Department of Health Promotion and Disease Prevention, Florida International University, Miami, FL, USA; Department of Psychology, The University of Texas at San Antonio, San Antonio, TX, USA

**Keywords:** adolescence, ethnic identity, fMRI, resting state functional connectivity, discrimination, ABCD Study

## Abstract

Ethnic identity refers to how individuals perceive and experience themselves in the context of social groups, racial background, or culture (Phinney & Ong., 2007). Ethnic identity is positively associated with psychological well-being (Rivas-Drake et al., 2014) and negatively associated with depression and anxiety (Forstmeier et al., 2021). Those with strong ethnic identity may display resiliency to the negative effects of discrimination on psychological well-being (Urzúa et al., 2021). Phinney’s model describes four profiles for how people put effort into, participate in, and reflect upon their ethnic identity (Phinney, 1989). Despite prior work addressing ethnic identity and psychosocial outcomes (for review, see Meca et al., 2023), few studies have considered its neurobiological underpinnings. In the current study, we identified profiles of ethnic identity among participants in The Adolescent Brain Cognitive Development Study (ABCD Study) using latent profile analysis. Next, we examined resting state functional connectivity differences across observed profiles and assessed the moderating effects of perceived discrimination. Results indicated heightened cingulo-parietal (CPAR) network connectivity among adolescents with highly diffuse ethnic identities; among moderately achieved ethnic identities, perceived discrimination moderated the association between ethnic identity and CPAR connectivity. We discuss how these findings may be related to attentional shift, error monitoring, autobiographical memory, and social judgements.

## Introduction

Ethnic identity is a sociocultural construct that refers to how individuals perceive, understand, and experience themselves in the context of a specific social group, racial background, or culture (Phinney and Ong, 2007). Ethnic identity is positively associated with adaptive psychosocial functioning, such as well-being, self-esteem, and coping behaviors (Rivas-Drake et al., 2014; Smith and Silva, 2011; Bracey et al., 2004; Turnage, 2004; Roberts et al., 1999) and negatively associated with maladaptive functioning, such as depression, anxiety, and substance use (Forstmeier et al., 2021; Fisher et al., 2017; Burnett-Zeigler et al., 2013; Williams et al., 2012; Roberts et al., 1999). Discrimination-related experiences play a key role in the relation between ethnic identity and psychosocial functioning (Nguyen et al., 2015; Romero et al., 2014; Street et al., 2008; Greene et al., 2006). Specifically, individuals with a strong ethnic identity are more resilient to the negative effects of discrimination on psychological well-being, as they possess a strong sense of belonging and pride in their ethnic group (Urzúa et al., 2021; Romero et al., 2014).

A widely known model proposes that ethnic identity is bidimensional (Roberts et al., 1999; Phinney, 1992),and includes a two-step process of first seeking knowledge about one’s ethnic identity (i.e., “exploration”) and then engaging in introspective work centered around one’s own ethnic identity, leading to a sense of belongingness and pride (i.e., “commitment”) (Mills and Murray, 2017; Phinney and Ong, 2007). Discrete categorizations, or profiles, of ethnic cultural exploration and commitment, or how people put effort into, participate in, and reflect upon their ethnicity can be identified by analyzing data-driven patterns across individuals (Cheon et al., 2020; Sanchez et al., 2016; Seaton et al., 2006). Phinney’s model proposes four profiles of ethnic identity: 1) **diffuse** (i.e., little to no exploration of or commitment to ethnic identity), 2) **foreclosed** (i.e., little to no exploration of, but commitment to ethnic identity), 3) **moratorium** (i.e., engaged exploration but no commitment to ethnic identity), and 4) **achieved** (i.e., engaged exploration resulting in commitment to ethnic identity) (**Figure 1**) (Seaton et al., 2006; Phinney, 1989).

**Figure 1.**
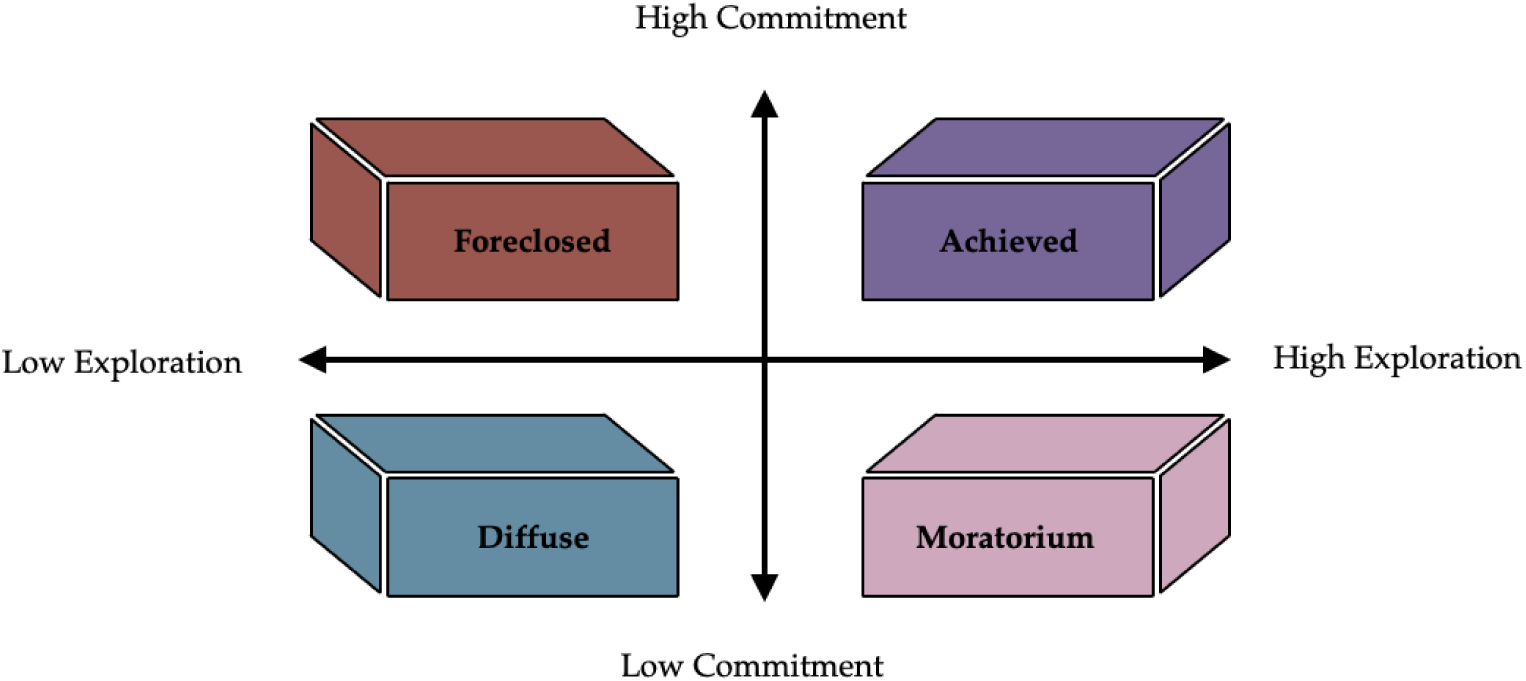
Phinney’s Model of Ethnic Identity. Each profile within Phinney’s model is characterized by dimensions of high or low exploration and high or low commitment.

Despite prior work addressing ethnic identity and psychosocial outcomes (for review, see Meca et al., 2023), few studies have considered neurobiological functioning. Among these, a significant study recently investigated the neural underpinnings of ethnic identity, revealing that brain networks associated with social functioning and cognitive control are linked to ethnic identity exploration and commitment (Constante et al., 2023). Using resting state functional connectivity (rsFC), Constant and colleagues found that default mode network (DMN) connectivity was positively associated with ethnic identity exploration, whereas higher frontoparietal network (FPN) connectivity was associated with ethnic identity commitment^1^(Constante et al., 2023). While these results provide important new insight into how ethnic identity may be related to functional brain organization, the authors’ continuous variable approach focused on exploration and commitment as separate constructs and did not consider them from a more holistic perspective. Moreover, despite the known associations between racial-ethnic discrimination, ethnic identity, and psychosocial functioning (Nguyen et al., 2015; Romero et al., 2014; Street et al., 2008; Greene et al., 2006), ethnic identity and rsFC models have not included discrimination experiences. Taken together, using a person-based framework to evaluate Phinney’s model may be helpful in examining rsFC differences across ethnic identity profiles. This would allow researchers to better understand fundamental mechanisms of identity processing, such as social cognition, which can help tailor strategies that promote adaptive psychosocial functioning or reduce maladaptive psychosocial functioning (Telzer et al., 2018).

In the current study, our overall objective was to examine rsFC differences across Phinney’s model of ethnic identity profiles. We first aimed to identify ethnic identity profiles using a person-centered approach within a large, demographically diverse sample from **The Adolescent Brain Cognitive Development**℠ **Study (ABCD Study®)** (Volkow et al., 2018). A limited amount of research exists on profiles of ethnic identity (Wantchekon & Umana-Taylor, 2021; Cheon et al., 2020); previous research has yielded mixed findings when considering various cultures, countries of origin, race, and ethnicity (Xie et al., 2021; Seaton et al., 2006; Phinney, 1989; Clark et al., 1976). Our goal was to validate and extend previous findings and uncover possible variations of ethnic identity subtypes using the latent profile analysis (LPA) method. We hypothesized that LPA of ethnic identity data from adolescent participants in the **ABCD Study®** would yield all four ethnic identity profiles described by Phinney’s model: i) diffuse, ii) foreclosed, iii) moratorium, and iv) achieved (Seaton et al., 2006; Phinney et al., 1998). After we identified LPA-derived adolescent ethnic identity profiles, we leveraged these results to explore profile membership effects on between- and within-network rsFC. Specifically, we expected DMN and salience network (SN) connectivity would be more strongly correlated with engaged exploration of ethnic identity (e.g., moratorium and achieved profiles) than diffuse ethnic identity, given their roles during self-referential processing and adaptive social behaviors (Webb et al., 2022; Marstaller et al., 2020; Holtz et al., 2012; Menon and Uddin, 2010), while DMN-FPN connectivity would be strongly correlated with commitment to ethnic identity (e.g., foreclosed and achieved profiles). Lastly, we conducted an exploratory analysis to assess the moderating effects of adolescent-reported perceived discrimination on identified ethnic identity profiles and rsFC. Finally, we discuss how investigating ethnic identity, rsFC, and perceived discrimination provides a deeper and more culturally focused contextualization of the social determinants of health on brain function and development.

## Methods

We utilized existing questionnaire and rsFC data from the Adolescent Brain Cognitive Development (ABCD) Study. Below we detailed the ABCD Study design, participants, measures, and rsFC data acquisition procedures. The analysis plan was pre-registered on Open Science Framework (https://osf.io/p4hkr).

### Participants

The ABCD Study is the largest longitudinal study of adolescent brain development and health in the United States (US) (Volkow et al., 2018). Across 21 US locations, ABCD Study applied an epidemiologically-based strategy to recruit a sample of geographically, demographically, and socioeconomically diverse adolescents and their caregivers (Compton et al., 2019). Each ABCD site had institutional review board (IRB) approval, and all adolescent participants provided informed assent to participate while caregivers gave informed consent. Extensive description regarding ABCD recruitment and assessment procedures are available (Garavan et al., 2018). Data from the ABCD Study were accessed via the NIMH Data Archive (NDA; https://nda.nih.gov) and the current study analyzed data from the ABCD Curated Annual Release 5.1. To identify ethnic identity profiles we used data from the Multiracial Ethnic Identity Measure (MEIM-R) at the Year 3 Follow-Up (Y3; 2019-2021), ages 12-13. Next, we tested for rsFC differences across profiles among youth who completed an in-person magnetic resonance imaging (MRI) session at the Year 4 Follow-Up (Y4; 2020-2022) at ages 13-14 where only half the data was available at the time of this study. Finally, we examined the moderating effects of perceived discrimination among adolescents, ages 13-14, who completed the Perceived Discrimination Scale at the Year 4 Follow-Up (Y4; 2020-2022).

### Measures

#### Participant Demographics

Adolescent and caregiver demographic variables were surveyed at baseline for the ABCD Study (2016-2018). Caregiver-reported adolescent demographics included age, gender identity, race, ethnicity, and country of origin. Caregiver-reported caregiver demographics included age, gender identity, race, ethnicity, education level, family income, and country of origin (Barch et al., 2018).

#### Multiracial Ethnic Identity Measure

To investigate profiles of ethnic identity, we utilized the MEIM-R self-report questionnaire that was administered as part of the ABCD Study. The MEIM-R is used to investigate ethnic identity status through assessment of efforts, participation, and reflection of one’s own ethnic cultural group. This included a 6-item assessment, which yields two factors: ethnic identity exploration (i.e., “*I have often talked to other people in order to learn more about my ethnic group*”) and commitment (i.e., “*I have a strong sense of belonging to my own ethnic group*”). The internal consistency was deemed ‘good’ according to alpha values (exploration α = 0.83, commitment α = 0.89) (Phinney & Ong, 2007).

#### Neuroimaging Data Acquisition and Preprocessing

Adolescent participants in the ABCD Study completed a neuroimaging protocol that included resting state functional MRI (fMRI) at Y4. FMRI data were acquired using high spatial and temporal resolution simultaneous multislice/multiband echo-planar imaging (EPI) (Hagler et al., 2019). For Siemens scanners, fMRI scan parameters were 90×90 matrix, 60 slices, field of view=216×216, echo time/repetition time=30/800ms, flip angle=52°, 2.4mm isotropic resolution, and slice acceleration factor 6. The complete protocols for all vendors and sequences are provided by Casey and colleagues (Casey et al., 2018). Participants were scanned while they completed four 5-minute resting state BOLD fMRI scans with their eyes open and fixated on a crosshair.

Imaging data preprocessing was performed by the ABCD Data Analysis, Informatics, and Resource Center (DAIRC; (Hagler et al., 2019). Data preprocessing included removal of primary frames, temporal filtering, and calculation of regions of interest (ROI). Measures of rsFC were generated using a cortical surface seed-based correlation approach. In the current study, tabulated rsFC data were analyzed from ROIs defined by the Gordon parcellation (Gordon et al., 2016). Averaged signal time series from all voxels within an ROI were used to generate Pearson’s correlation values between pairs of ROIs. These correlations were then transformed into normally distributed Fisher z-values to assess between- and within-network connectivity strength. Within-network correlations were computed by taking the mean of the Fisher-transformed correlations for each of the pairwise ROIs in a given Gordon network. Between-network correlations were computed by taking the mean of ROIs from two Gordon networks (Hagler et al., 2019).

#### Measure of Perceived Discrimination

ABCD participants also completed the Measure of Perceived Discrimination (MPD), a 7-item subscale from the self-report questionnaire of Perceived Discrimination Scale. The MPD had an acceptable internal consistency (α = 0.81) and investigated the feelings, experiences, and frequency of unjust treatment due to ethnicity. An example from this subset of questions is: “*How often do the following people treat you unfairly or negatively because of your ethnic background*?” (Phinney et al., 1998). For the current study, we used the summary score that reflects the mean responses across items (Gonzalez et al., 2021).

### Analyses

#### Latent Profile Analysis

We performed an LPA of MEIM-R responses to understand ethnic identity from a person-centered approach. LPA is a statistical method that aggregates patterns of participant responses into data-driven groupings. In the current study, we conducted an LPA to identify groups of adolescent participants with similar ethnic identity profiles. LPA was performed in R using the tidyLPA package (Rosenberg et al., 2018). Model fit indices included Akaike’s Information Criteria (AIC), Bayesian Information Criteria (BIC), entropy, and bootstrapped likelihood ratio test (LRT). AIC and BIC were assessed to understand goodness of fit and probable number of models. The lowest AIC and BIC values in the model are optimal. Entropy was utilized to evaluate certainty of participant profile classification, values greater than or equal to 0.8 are optimal. LRT was computed to identify p-values and fit improvement of proposed models. Significant LRT p-values indicated the additional profile was a significant fit improvement and should be used in the final analysis relative to *k*-1 profiles. To visualize data grouping results at 95% confidence, the tidyLPA plot_profile was used.

To contextualize these profiles, *X*^2^ and *t*-tests were conducted to assess significant demographic differences across ethnic identity. Demographic variables of interest included: adolescent age, gender identity, race, ethnicity, and country of origin, as well as caregiver age, gender identity, race, ethnicity, education level, family income, and country of origin.

#### Resting State Functional Connectivity Analyses

To understand how ethnic identity profiles may be associated with differences in rsFC we examined the between- and within-network connectivity correlation matrix for eight Gordon network parcels. The parcels included were: Cingulo-Opercular (CON), Cingulo-Parietal (CPAR), Default Mode (DMN), Dorsal Attention (DAN), Fronto-Parietal (FPN), Retrosplenial-Temporal (RT), Salience (SN), and Ventral Attention (VAN). These specific parcels represented a subset of the original 13 Gordon networks in which the higher-order cognitive parcels of interest were included while the primary sensory (i.e., Auditory, Visual) and sensorimotor (i.e., Sensorimotor-Hand and Sensorimotor-Mouth) parcels were excluded (“None” was also excluded). Covariates included adolescent age, gender identity, race, ethnicity, as well as caregiver age, education, and family income. Linear mixed-effect models (LMM) methods were utilized as these are robust for unequal groups of data, controlling covariates, and hierarchical and longitudinal datasets (Dick et al., 2021). Random effects for site and family were constructed within the model to account for the nested structure of ABCD data (Saragosa-Harris et al., 2022). We applied the Benjamini-Hochberg correction to control for the false discovery rate (FDR) at 0.05 p-value significance due to multiple comparisons (Benjamini and Hochberg, 1995). Missing data were handled via casewise deletion.

#### Moderation of Perceived Discrimination

To further explain the relation between ethnic identity profiles and rsFC, we examined the MPD measure using LMM (**Figure 2**) and corresponding random effects modeling as constructed in the previous analysis. Here, the use of LMM was selected to maintain consistency across analyses and provide a robust framework for assessing moderation (Brown, 2021). Covariates included adolescent age, gender identity, and race, ethnicity, as well as caregiver age, education, and family income. Again, we applied the Benjamini-Hochberg correction (Benjamini and Hochberg, 1995) in the same manner as the previous analysis.

**Figure 2.**
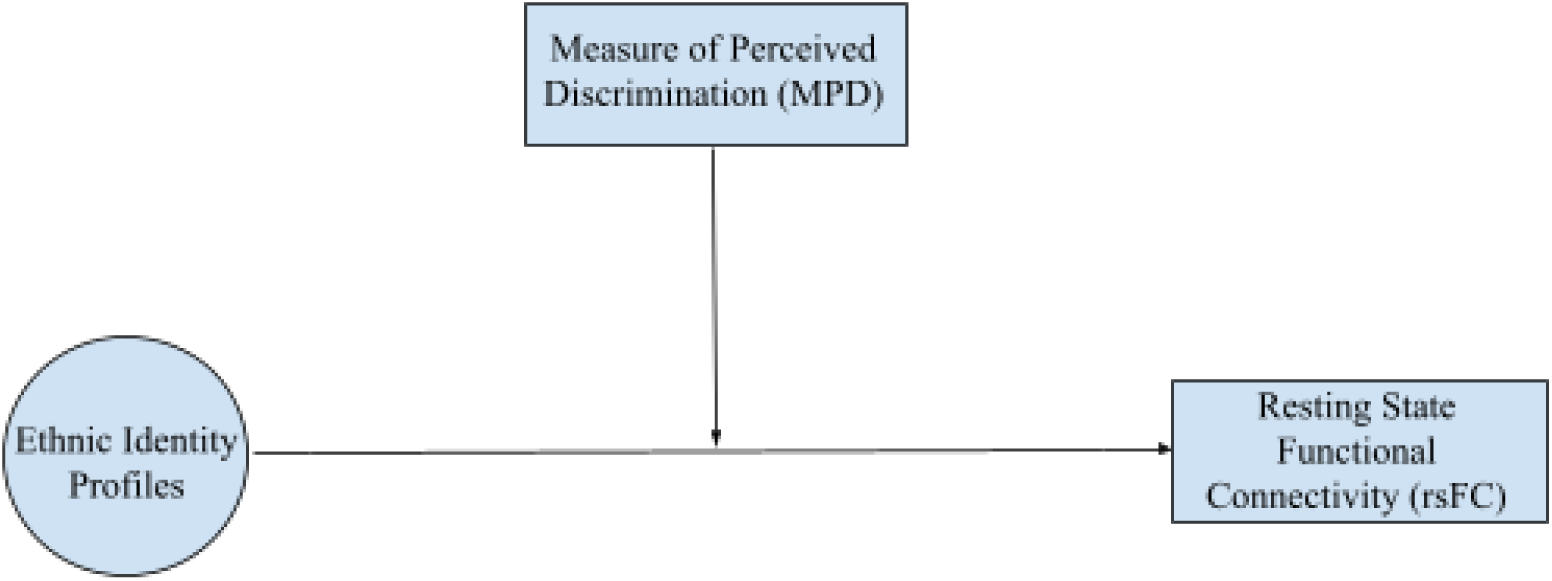
LMM Statistical Framework and Moderation of Perceived Discrimination. An LMM approach was used for our exploratory moderation analysis. We assessed how perceived discrimination impacts the relation between ethnic identity profiles and rsFC. Demographic variables of interest were added as covariates.

## Results

### Participants

LPA was conducted on all ABCD participants who completed the MEIM-R at Y3, consisting of 8,164 adolescents (*N* = 8,164) (**Table 1**). rsFC differences were tested among a subset of those participants who completed the neuroimaging session at Y4, reducing the sample size to 2,560 adolescents (*N* = 2,560). Finally, the moderation analysis was performed on a subset of participants who completed the MPD at Y4, which included 2,524 youth (*N* = 2,524). Demographic differences across samples revealed adolescent age (*F* = 7.909, *df* = 2, *p* < 0.004), adolescent race (*X^2^* = 52.132, *df* = 32, *p* = 0.014), and caregiver race (*X^2^* = 49.20, *df* = 34, *p* = 0.044) were significant (**Table 2**).

**Table 1.**
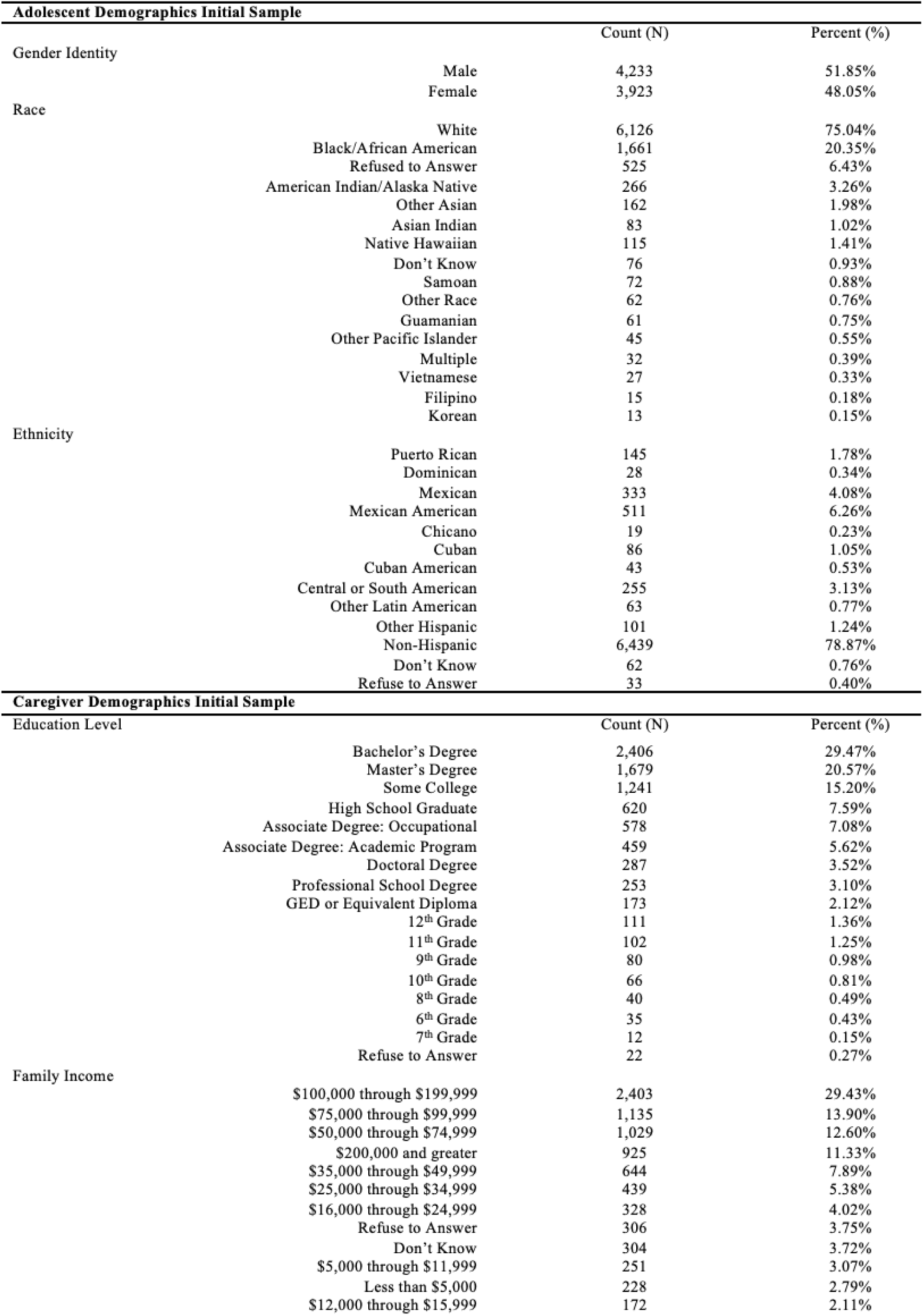
Demographics of MEIM-R Y3 Sample. The full demographic breakdown is provided in Supplement Table S1.

**Table 2.** Demographic Differences Between Samples. Statistical differences among the MEIM-R LPA sample (*N* = 8,164), resting-state functional connectivity (rsFC) sample (*N* = 2,560), and measure of perceived discrimination sample (*N* = 2,524) were examined. Degrees of freedom are denoted as df.

|  | df | Statistic | p-value |
| --- | --- | --- | --- |
| **Adolescent Demographics** |  |  |  |
| Age | 2 | $F = 7.909$ | <0.005 |
| Gender Identity | 14 | $X^2 = 3.448$ | 0.998 |
| Race | 32 | $X^2 = 52.132$ | 0.014 |
| Ethnicity | 24 | $X^2 = 16.389$ | 0.874 |
| Country of Origin | 134 | $X^2 = 57.807$ | 0.999 |
| **Caregiver Demographics** |  |  |  |
| Age | 2 | $F = 2.681$ | 0.069 |
| Gender Identity | 12 | $X^2 = 4.385$ | 0.975 |
| Race | 34 | $X^2 = 49.20$ | 0.044 |
| Ethnicity | 22 | $X^2 = 19.854$ | 0.592 |
| Education Level | 40 | $X^2 = 35.131$ | 0.689 |
| Family Income | 22 | $X^2 = 21.250$ | 0.505 |
| Country of Origin | 224 | $X^2 = 97.278$ | 0.999 |

### Latent Profile Analysis

Fit indices identified the model with five profiles to have the lowest Bayesian Information Criteria (BIC = 35769.33) and lowest Akaike Information Criterion (AIC = 35657.21), suggesting five profiles was a more optimal fit than four profiles (**Table 3**). In further support of BIC interpretation, results of the bootstrapped LRT p-values revealed significant differences in improvement of fit between models with four and five profiles (*p* = 0.01). Improvement of fit was also shown for differences of the three- and four-profile models (*p* = 0.01), two- and three-profile models (*p* = 0.01), and one- and two-profile models (*p* = 0.01). Probability minimum and maximum indicated the probability of each participant falling into a profile was fairly high. Regarding entropy, a predetermined cut-off value greater than or equal to 0.8 was deemed as a good fit for remaining as a distinct classification. The current results showed no models had an entropy value greater than 0.8; the five-profile model had the highest entropy at 0.79. Although an entropy of 0.8 was the pre-determined cut-off value, entropy interpretation alone should not determine the best fitting model. In some cases, models with a lower entropy but optimal values of BIC and bootstrapped LRT can still be considered a viable model (Sinha et al., 2021). Thus, a five-profile model was deemed optimal considering BIC, entropy, bootstrapped LRT, and theoretical interpretation of classes.

**Table 3.** Latent Profile Analysis Models.

| Classes | AIC | BIC | Entropy | N<br>Minimum | N<br>Maximum | Probability<br>Minimum | Probability<br>Maximum | LRT p-value |
| --- | --- | --- | --- | --- | --- | --- | --- | --- |
| 1 | 40148.25 | 40176.28 | 1.00 | 100% | 100% | 1.00 | 1.00 | 1.00 |
| 2 | 37510.86 | 37559.92 | 0.61 | 41% | 59% | 0.86 | 0.89 | 0.01 |
| 3 | 36422.25 | 36492.32 | 0.68 | 15% | 61% | 0.79 | 0.89 | 0.01 |
| 4 | 35951.15 | 36042.25 | 0.72 | 4% | 54% | 0.71 | 0.88 | 0.01 |
| 5 | 35657.21 | 35769.33 | 0.79 | 4% | 39% | 0.81 | 0.90 | 0.01 |

After attaining each participant’s profile membership, we utilized posterior probabilities greater than or equal to 0.7 to ensure profile stability. After posterior probability was determined, our sample size was reduced to N = 6,923. These five profiles were compared to the profiles described by Phinney’s model and cross-validated by visualizing participant z-scores for each profile dimension. Following this procedure, the data-driven profiles were defined as: 1) **highly diffuse** (i.e., very low scores of little to no exploration of or commitment to ethnic identity), 2) **moderately diffuse** (i.e., moderately low scores of little to no exploration of or commitment to ethnic identity), 3) **foreclosed** (i.e., little to no exploration of, but commitment to ethnic identity), 4) **moderately achieved** (i.e., moderately engaged exploration resulting in commitment to ethnic identity), 5) **highly achieved** (i.e., extremely engaged exploration resulting in commitment to ethnic identity) (**Figure 3**). We found a significant difference in profile membership among caregiver age; no other significant differences in profile membership were observed for all other adolescent and caregiver demographics (**Table 4**).

**Figure 3.**
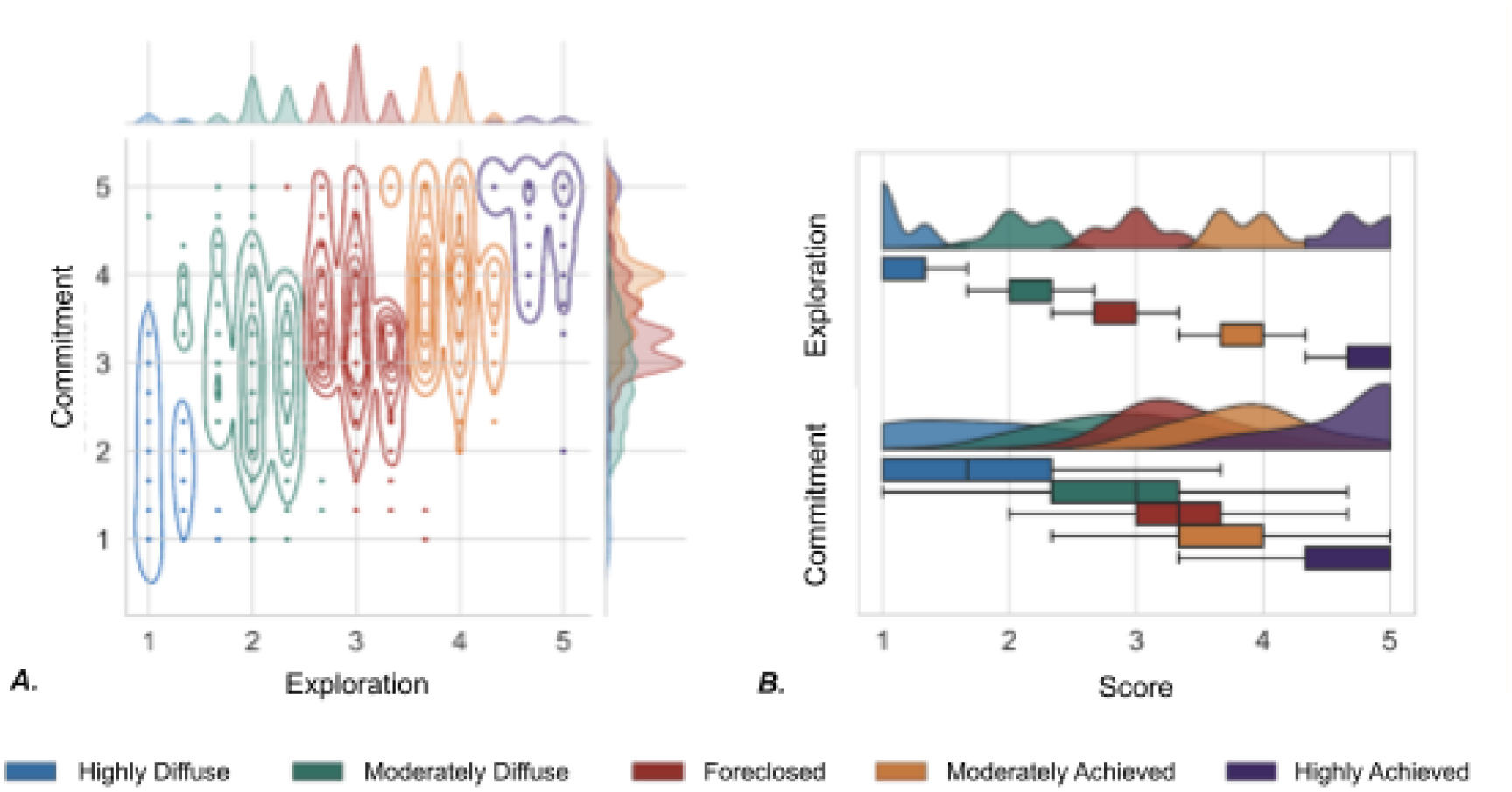
Ethnic Identity Exploration and Commitment Across Profiles. Latent profile analysis revealed five ethnic identity profiles among adolescents in the ABCD Study. Ethnic identity subscales for exploration and commitment are shown in (***A***) a joint kernel density estimate plot and (***B***) a raincloud plot. In both visualizations, participants in the highly diffuse profile (blue) exhibited lowest values for exploration and commitment, moderately diffuse participants (green) were moderately low on exploration and commitment, foreclosed participants (red) were high on commitment but low on exploration, moderately achieved participants (orange) were somewhat high on both exploration and commitment, and highly achieved participants (purple) were observed to have the highest values for both exploration and commitment.

**Table 4.** Adolescent and Caregiver Demographic Differences among Profiles. Variables which contained less than or equal to 5 participants were collapsed into reported categories to maintain anonymity (CPRD, 2024).

|  |  | Highly Diffuse<br>N = 2529 | Moderately Diffuse<br>N = 1753 | Foreclosed<br>N = 277 | Moderately<br>Achieved<br>N = 370 | Highly<br>Achieved<br>N = 1994 | Statistic | p-value |
| --- | --- | --- | --- | --- | --- | --- | --- | --- |
| **Adolescent Demographics** |  |  |  |  |  |  |  |  |
| Age M(SD)) | | 9.96 (0.63) | 9.93 (0.62) | 9.91(0.63) | 0.92 (0.62) | 9.97 (0.62) | $F = 1.736$ | 0.240 |
| Gender Identity (N) | | | | | | | $\chi^2 = 34.461$ | 0.251 |
|  | Male | 1,305 | 947 | 169 | 172 | 977 |  |  |
|  | Female | 1,220 | 805 | 107 | 198 | 1,015 |  |  |
| Race (N) | | | | | | | $\chi^2 = 61.663$ | 0.290 |
|  | White | 1,990 | 1,537 | 244 | 179 | 1,292 |  |  |
|  | Black/African American | 418 | 187 | 30 | 165 | 577 |  |  |
|  | American Indian/Native American | 82 | 41 | 8 | 18 | 83 |  |  |
|  | Other | 376 | 178 | 32 | 75 | 413 |  |  |
| Ethnicity (N) | | | | | | | $\chi^2 = 39.897$ | 0.632 |
|  | Hispanic | 580 | 402 | 65 | 91 | 467 |  |  |
|  | Non-Hispanic | 1,949 | 1,351 | 212 | 279 | 1,527 |  |  |
| Country of Origin (N) | | | | | | | $\chi^2 = 214.190$ | 0.445 |
|  | Born in US | 2,347 | 1,650 | 257 | 335 | 1,842 |  |  |
|  | Born Outside US | 182 | 103 | 20 | 35 | 152 |  |  |
| **Caregiver Demographics** |  |  |  |  |  |  |  |  |
| Age (M(SD)) | | 40.2 (7.9) | 40.4 (8.2) | 41.0 (7.7) | 41.6 (7.8) | 41.1 (7.9) | $F = 15.58$ | <0.001 |
| Gender Identity (N) | | | | | | | $\chi^2 = 21.383$ | 0.549 |
|  | Male | 654 | 431 | 63 | 96 | 462 |  |  |
|  | Female | 1,875 | 1,322 | 214 | 274 | 1,532 |  |  |
| Race (N) | | | | | | | $\chi^2 = 73.871$ | 0.134 |
|  | White | 1,954 | 1,530 | 238 | 175 | 1,262 |  |  |
|  | Black/African American | 334 | 127 | 22 | 145 | 468 |  |  |
|  | American Indian/Native American | 57 | 32 | 6 | 16 | 58 |  |  |
|  | Other | 303 | 121 | 22 | 59 | 350 |  |  |
| Ethnicity (N) | | | | | | | $\chi^2 = 35.092$ | 0.681 |
|  | Hispanic | 494 | 345 | 44 | 69 | 341 |  |  |
|  | Non-Hispanic | 2,035 | 1,408 | 233 | 301 | 1,653 |  |  |
| Education Level (N) | | | | | | | $\chi^2 = 62.062$ | 0.925 |
|  | Less than HS | 289 | 221 | 32 | 47 | 243 |  |  |
|  | HS/GED | 619 | 465 | 79 | 98 | 552 |  |  |
|  | Some College | 738 | 520 | 93 | 122 | 678 |  |  |
|  | College Grad or Higher | 883 | 547 | 73 | 103 | 521 |  |  |
| Family Income (N) | | | | | | | $\chi^2 = 55.694$ | 0.128 |
| | < \$50k | 679 | 438 | 80 | 95 | 473 | | |
| | \$50k-\$100k | 674 | 503 | 75 | 109 | 563 | | |
| | >\$100k | 1,176 | 812 | 122 | 166 | 958 | | |
| Country of Origin (N) | | | | | | | $\chi^2 = 409.39$ | 0.456 |
|  | Born in US | 2,325 | 1,603 | 255 | 342 | 1,851 |  |  |
|  | Born Outside US | 204 | 150 | 22 | 28 | 143 |  |  |

### Resting State Functional Connectivity

To examine differences in rsFC across ethnic identity profiles, we used an LMM approach on the between- and within-network correlation matrix for eight Gordon network parcels. Results indicated that the highly diffuse ethnic identity profile was positively associated with within-cingulo-parietal network (CPAR) connectivity (*β* = 0.110, *SE* = 0.038, *t*(1435.260) = 2.910, *p* = 0.004, *p*_FDR_ = 0.026) (**Figure 4**). Significant covariates include those who reported Mexican American ethnicity (*β* = −0.117, *SE* = 0.040, *t*(1591.102) = −2.892, *p* = 0.004, *p*_FDR_ = 0.026), Other Hispanic ethnicities (*β* = −0.177, *SE* = 0.054, *t*(1683.289) = −3.25, *p* = 0.001, *p*_FDR_ = 0.020), Black/African American race (*β* = −0.060, *SE* = 0.016, *t*(1581.008) = −3.81, *p* = 0.001, *p*_FDR_ = 0.005), and caregiver income (*β* = 0.008, *SE* = 0.003, *t*(1712.219) = 2.96, *p* = 0.003, *p*_FDR_ = 0.026). No other significant results survived the correction for multiple comparisons (Table S2).

**Figure 4.**
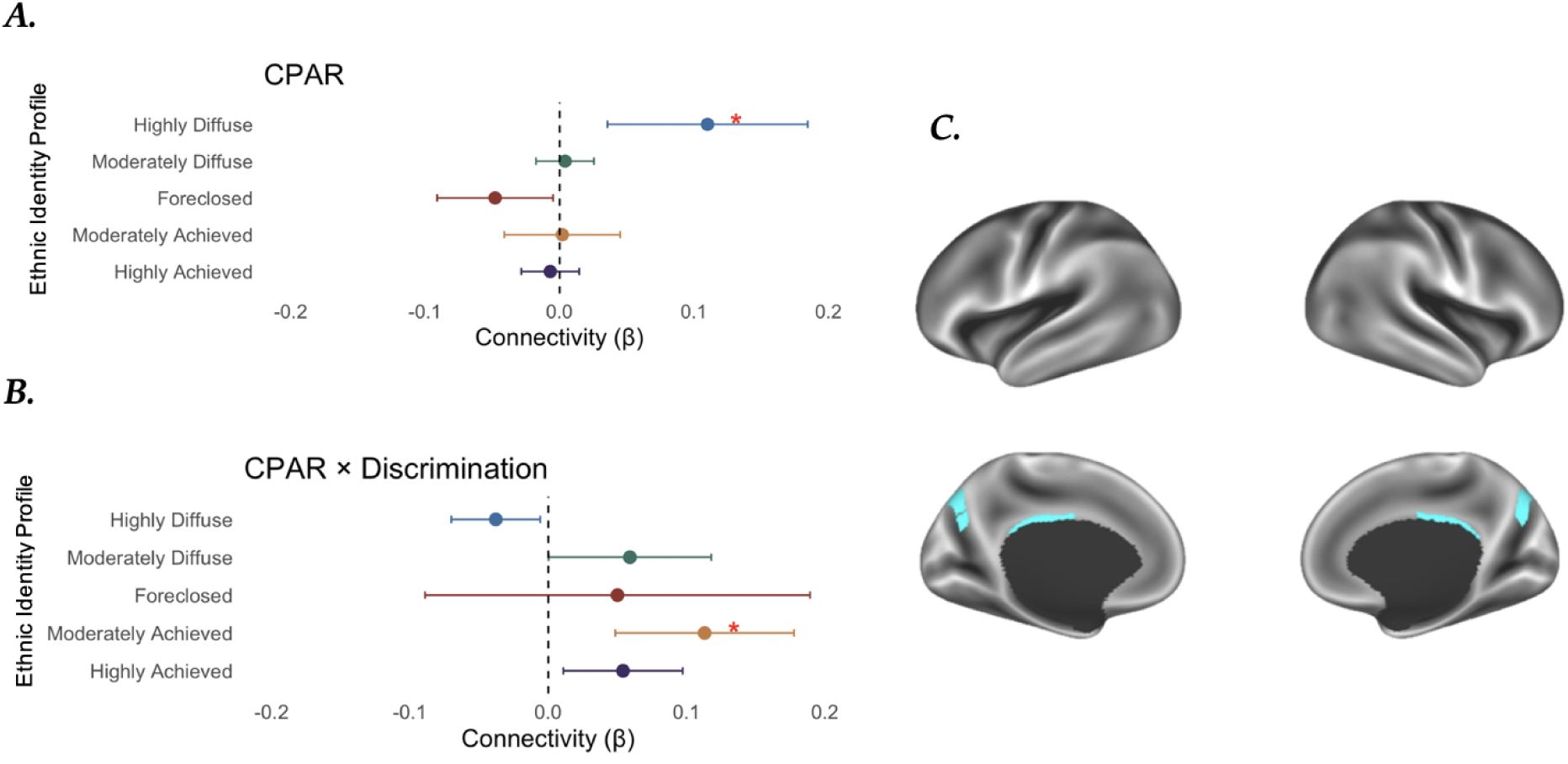
Within CPAR and Discrimination Interaction Results. Forest plots of within-network connectivity (beta estimates) for each profile for (***A***) Cingulo-Parietal (CPAR) and (***B***) CPAR x Discrimination, highlighting the unit-change, intensity, precision, and directionality of findings. (C) maps of the CPAR network according to the Gordon parcellation (Van Essen et al., 2017; Gordon et al., 2016).

### Moderation of Perceived Discrimination

LMM results indicated that among the moderately achieved participants, perceived discrimination moderated the association between ethnic identity profile and within-CPAR connectivity (*β* = 0.113, *SE* = 0.033, *t*(1915.73) = 3.42, *p* = 0.0006, *p*_FDR_ = 0.012) (**Figure 4**). Significant covariates include those who reported Mexican American (*β* = −0.120, *SE* = 0.0340, *t*(1593.168) = −2.97, *p* = 0.003, *p*_FDR_ = 0.023), Other Hispanic ethnicities (*β* = −0.179, *SE* = 0.054, *t*(1686.04) = −3.29, *p* = 0.001, *p*_FDR_ = 0.013), Black/African American race (*β* = −0.066, *SE* = 0.016, *t*(1608.40) = −4.09, *p* = 0.001, p_FDR_ = 0.002), and caregiver income (*β* = 0.007, *SE* = 0.003, *t*(1713.06) = 2.69, *p* = 0.007, *p*_FDR_ = 0.046). No other significant finding survived the correction for multiple comparisons (Table S3).

## Discussion

We applied a person-centered, multimodal approach, including self-report and brain-based measures, to better understand ethnic identity and resting state functional connectivity (rsFC), with the goal of gaining deeper cultural insights into the role of the social determinants on health and brain function. To this end, we examined associations between ethnic identity, rsFC, and perceived discrimination among adolescents using data from the ABCD Study. LPA revealed five profiles of ethnic identity, including **highly diffuse** (i.e., very low to no exploration of or commitment to ethnic identity), **moderately diffuse** (i.e., moderately low exploration of or commitment to ethnic identity), **foreclosed** (i.e., low to no exploration of, but commitment to ethnic identity), **moderately achieved** (i.e., moderately engaged exploration resulting in commitment to ethnic identity), and **highly achieved** (i.e., extremely engaged exploration resulting in commitment to ethnic identity). In terms of rsFC, the highly diffuse ethnic identity profile was significantly associated with within-CPAR connectivity. Furthermore, participants in the moderately achieved profile who reported higher levels of perceived discrimination exhibited increased within-CPAR connectivity.

### Ethnic Identity Profiles Among Adolescents in the ABCD Study

The ABCD Study is the largest longitudinal study of brain and health development to date. To our knowledge, the current study examined the largest and most diverse sample of adolescents using a person-centered, data-driven approach to ethnic identity. Our findings suggest adolescents ages 12-13 experience efforts, participation, and reflection of ethnic identity at differing intensity. In the ABCD sample, diffuse ethnic identity was found to be categorized by high and moderate levels via an extremely low mean z-score for exploration and commitment (i.e., highly diffuse) and a moderately low z-score for exploration and commitment (i.e., moderately diffuse). Thus, those within the highly diffuse profile may experience little to no efforts, participation, and reflection of ethnic identity, whereas those within the moderately diffuse ethnic identity may experience mild efforts, participation, and reflection of ethnic identity. Achieved ethnic identity was also found to be categorized by high and moderate levels. Similar to diffuse profiles, we identified an extremely high mean z-score for exploration and commitment (i.e., highly achieved) and a moderately high z-score for exploration and commitment (i.e., moderately achieved). Thus, a similar pattern may also be true for achieved ethnic identity where those within the highly achieved profile have the highest reports of effort, participation, and reflection of ethnic identity and those within the moderately achieved ethnic identity have moderate reports of effort, participation, and reflection. Notably, previous research has identified both achieved and highly achieved profiles, parallel with our findings, among Black Americans (Driscoll et al., 2024). Finally, while our results are broadly aligned with Phinney’s theoretical model, we note that no moratorium profile was identified; that is, no adolescents were identified as being engaged in exploration but without commitment to their ethnic identity. This suggests that adolescents ages 12-13 may not be engaged in seeking new information about their ethnic identity unless there is already some level of commitment. This is supported by past evidence suggesting adolescents are less cognitively flexible and less self reflective compared to emerging adults, which may result in higher sensitivity to social evaluation, difficulties of emotional regulation, and outcomes of anxiety and depression (Gao et al., 2025; Ruan et al., 2023; Blakemore and Mills, 2014; Umaña-Taylor et al., 2014; Silvers et al., 2012). Overall, our findings are mostly consistent with Phinney’s theorized model (Seaton et al., 2006; Phinney, 1989), and show strong correspondence to more recent empirical evidence demonstrating lack of moratorium findings and moderate levels of exploration and commitment (Maehler, 2022). In that study, diffuse (i.e., low exploration and commitment) and achieved (i.e., high exploration and commitment) profiles were observed in addition to marginally committed (i.e., moderately low exploration and commitment) and searching while committed (i.e., moderately high exploration and commitment) (Maehler, 2022). These map on closely to highly and moderately diffuse and achieved profile standardized scores found in the current study (Figure S1).

### CPAR Connectivity

Next, we investigated the relations of these five ethnic identity profiles with rsFC and perceived discrimination. We found an association between within-CPAR connectivity and the highly diffuse profile; in addition, we found that perceived discrimination moderated the association between within-CPAR connectivity and the moderately diffuse profile. Notably, our hypotheses of increased DMN-SN and DMN-FPN connectivity were not met; instead, our results emphasized CPAR regions, including the precuneus and posterior cingulate cortex, which are associated with attentional shift control, error monitoring, and risk avoidance (Corlett et al., 2022; Roy et al., 2011; Nagahama et al., 1999). Thus, it is possible that heightened attention and error monitoring mechanisms may act as a downstream avoidance response, much like what we see in vigilance–avoidance theory (Derakshan et al., 2007), among those within the highly diffuse profile by perceiving new information of ethnic identity as error, and therefore, allowing for disengagement of identity exploration and commitment processes. Such avoidance agrees with reports of lowest exploration and commitment wherein individuals do not have an established ethnic identity, and are not engaged in thinking about their ethnic identity (Seaton et al., 2006; Phinney, 1989). Furthermore, increased CPAR connectivity may help explain how adolescents experiencing highly diffuse ethnic identity commonly report maladaptive responses, such as depression and anxiety (Chavez-Korell and Torres, 2014; Yip et al., 2006). In addition to enhanced error monitoring and avoidance, self-referential thinking and internally focused attention are also associated with CPAR functioning and are commonly dysregulated in depression (Schreiner et al., 2019; Zhu et al., 2012). High rumination, a common symptom of depression, is also commonly linked to precuneus functioning (Cheng et al., 2024). Additionally, self-referential thinking and internal focus of attention may be associated with anxiety through decreases in decision-making processes (Moser et al., 2013). This follows previous literature on maladaptive exploration of ethnic identity that suggests individuals may experience indecisiveness when there are indications of high rumination (Luyckx et al., 2008). Taken together, the current CPAR connectivity findings may help explain the neural underpinnings of the associations between individuals with highly diffuse ethnic identity and psychosocial outcomes such as anxiety and depression.

Autobiographical memory and social judgements have also been implicated in CPAR functioning (Agathos et al., 2023; Wainberg et al., 2022; Cavanna and Trimble, 2006; Gilboa et al., 2004). Regarding the interaction of perceived discrimination and CPAR among moderately achieved individuals, these mechanisms align with findings that a more developed ethnic identity can buffer against discrimination (Yip et al., 2019). Specifically, among individuals with moderately achieved ethnic identity, self-appraisal in relation to others and ability to detect errors related to one’s own thought processes may be more evident when discrimination experiences are heightened. Previous evidence suggests achieved and positive ethnic identity profiles who face socio-structural and socio-individual barriers such as discrimination are more likely to respond adaptively (Romero et al., 2014; Galliher et al., 2011; Rivas-Drake et al., 2008), which may be better explained by these mechanisms related to heightened CPAR connectivity in the current results.

Additionally, social judgements within the family may also create a buffer of self-sustainability during discrimination for moderately achieved individuals. During achieved statuses of ethnic identity, understanding and accepting self-concepts related to cultural expectations from family and community are heavily assessed among individuals (Seaton et al., 2006; Phinney, 1989). These family narratives have the potential to buffer against discrimination while also influencing personal narratives within autobiographical memory (Weststrate et al., 2024; Camia et al., 2021). This integration of personal and cultural narratives reflected in achieved ethnic identity has the potential to impact positive well-being outcomes (Wang, 2021). For example, when examining culturally specific mechanisms of mother and child reminiscing, Chinese immigrant children experienced more adaptive behaviors when discussing negative emotions and cognitive coping with their mothers compared to European American children (Koh and Wang, 2021). This evidence along with the current findings highlight the importance of understanding cultural norms and family expectations as they relate to autobiographical memory to nurture adaptive outcomes and buffer against discrimination for individuals within the moderately achieved profile.

### Limitations

Several key limitations of our study and its findings should be noted. First, the MEIM-R was administered at the ABCD Year 3 Follow-Up and resting state fMRI data were collected at the ABCD Year 4 Follow-Up. Adolescents experience rapid developmental changes and a one-year difference in data collection could capture significantly different developmental stages. Second, the current study was also unable to replicate the moratorium ethnic identity profile that has been previously identified. Although there has been extensive literature suggesting adolescents experience low commitment and high exploration of ethnic identity, the lack of replication may be due to our large sample size; that is, the ABCD sample may present limitations for identifying nuances of profile categorization among subpopulations. Similarly, the attrition related to casewise deletion may have inadvertently caused subpopulations to be over- or underrepresented. Another limitation within the current study involves the use of resting state fMRI data. This method of assessing brain network connectivity does not directly elicit task-specific brain networks related to ethnic identity profiles (Zhang et al., 2016). However, the benefit to using rsFC is that we are able to examine several brain networks simultaneously. Additionally, the brain is also in an unconstrained, naturalistic state, making the results more interpretable than task-based results for ongoing experiences (Gonzalez-Castillo et al., 2021). Future task-based fMRI data would be beneficial as it would allow for probing of cognitive control, flexibility, and self-other processing. Finally, while the current study examined ethnic identity commitment and exploration as measured by the MEIM-R, it did not consider other dimensions such as positive and negative affect or centrality of ethnic identity. Such consideration may have resulted in differing solutions such as diffuse and low regard, diffuse and high regard, and developed and idealized, which have also been associated with adaptive psychosocial outcomes (Wantchekon & Umana-Taylor, 2021). Including these dimensions in future work could provide more granularity of neural underpinnings associated with psychological well-being among subpopulations.

## Conclusions

The current study identified five ethnic identity profiles among adolescents enrolled in the ABCD Study. Within-CPAR connectivity, linked to attentional shift, error monitoring, autobiographical memory, and social judgements, was associated with highly diffuse identities, while perceived discrimination moderated the association between within-CPAR connectivity and the moderately diffuse profile. Significant differences in caregiver age across profiles may warrant additional future analyses as such differences can be associated with parental identity and downstream effects of youth well-being (Fadjukoff et al., 2016). In addition, future work should investigate developmental patterns of ethnic identity, such as progression theory, which states that, in most cases, individuals experiencing early ethnic identity development during adolescence will initially endorse lower reports of ethnic identity (i.e., diffuse) and then move into achieved ethnic identity profiles (i.e., achieved) later into late adolescence (Phinney and Chavira, 1992; Phinney, 1989). In some cases, individuals progress from little to no exploration of or commitment to their ethnic identity (i.e., diffuse) to commitment without exploration (e.g., foreclosed) and then move to and stay in the foreclosed profile later in adolescence (Seaton et al., 2006). Identifying these differing patterns of progression from one ethnic identity profile to another, as well as the associated neurodevelopmental trajectories, could help clinicians better understand behaviors associated with specific trajectories, such as highly diffuse to foreclosed. Overall, our results suggest the need for longitudinal neurodevelopmental frameworks to account for reconsideration of ethnic identity and discrimination experiences when investigating adolescent ethnic identity trajectories.

## Acknowledgments

Primary funding for this project was provided by the FIU Embrace Center for Advancing Inclusive Communities. Data used in the preparation of this article were obtained from the Adolescent Brain Cognitive Development^SM^ (ABCD) Study (https://abcdstudy.org), held in the NIMH Data Archive (NDA). This is a multisite, longitudinal study designed to recruit more than 10,000 children aged 9-10 and follow them over 10 years into early adulthood. The ABCD Study® is supported by the National Institutes of Health and additional federal partners under award numbers U01DA041048, U01DA050989, U01DA051016, U01DA041022, U01DA051018, U01DA051037, U01DA050987, U01DA041174, U01DA041106, U01DA041117, U01DA041028, U01DA041134, U01DA050988, U01DA051039, U01DA041156, U01DA041025, U01DA041120, U01DA051038, U01DA041148, U01DA041093, U01DA041089, U24DA041123, U24DA041147. A full list of supporters is available at https://abcdstudy.org/federal-partners.html. A listing of participating sites and a complete listing of the study investigators can be found at https://abcdstudy.org/consortium_members/. ABCD consortium investigators designed and implemented the study and/or provided data but did not necessarily participate in the analysis or writing of this report. This manuscript reflects the views of the authors and may not reflect the opinions or views of the NIH or ABCD consortium investigators. For registered trademarks: “**ABCD Study®, Teen Brains. Today’s Science. Brighter Future.®, El cerebro adolescente. La ciencia de hoy. Un futuro más brillante.® and the ABCD Study Logos** are registered marks of the U.S. Department of Health & Human Services (HHS).” For service marks: “**Adolescent Brain Cognitive Development**℠ **Study, El Estudio del Desarrollo Cognitivo y Cerebral del Adolescente**℠, are service marks of the US Department of Health & Human Services (HHS).”

The ABCD data repository grows and changes over time. The ABCD data used in this report came from NDA ABCD Release 5.1 (DOI: 10.15154/1520591).

## Data and Code Availability

Code for this study can be found on Github (https://github.com/NBCLab/MEIM-ID-Profiles).

## Author Contributions

ARL, AM, and TRJ conceived and designed the project. TRJ and ML preprocessed data. TRJ analyzed data; TRJ, ARL, CH, ML, and AM interpreted data. DDS, JAP, and ML contributed scripts and pipelines. TRJ, ARL wrote the paper and all authors contributed to the revisions and approved the final version.

## Footnotes

1 Constante et al. refer to the dimensionality of ethnic identity using exploration and resolution where resolution refers to having clarity about one’s ethnic identity. Here we use the term commitment in place of resolution due to vast similarity of construct definition and to align with the corresponding ethnic identity measure used by the ABCD Study.

