## Supplement File for "Ethnic Identity Profiles Among Adolescents in the ABCD Study: Associations with Resting State Functional Connectivity and Perceived Discrimination"

### **Supplemental Results**

#### **Participants**

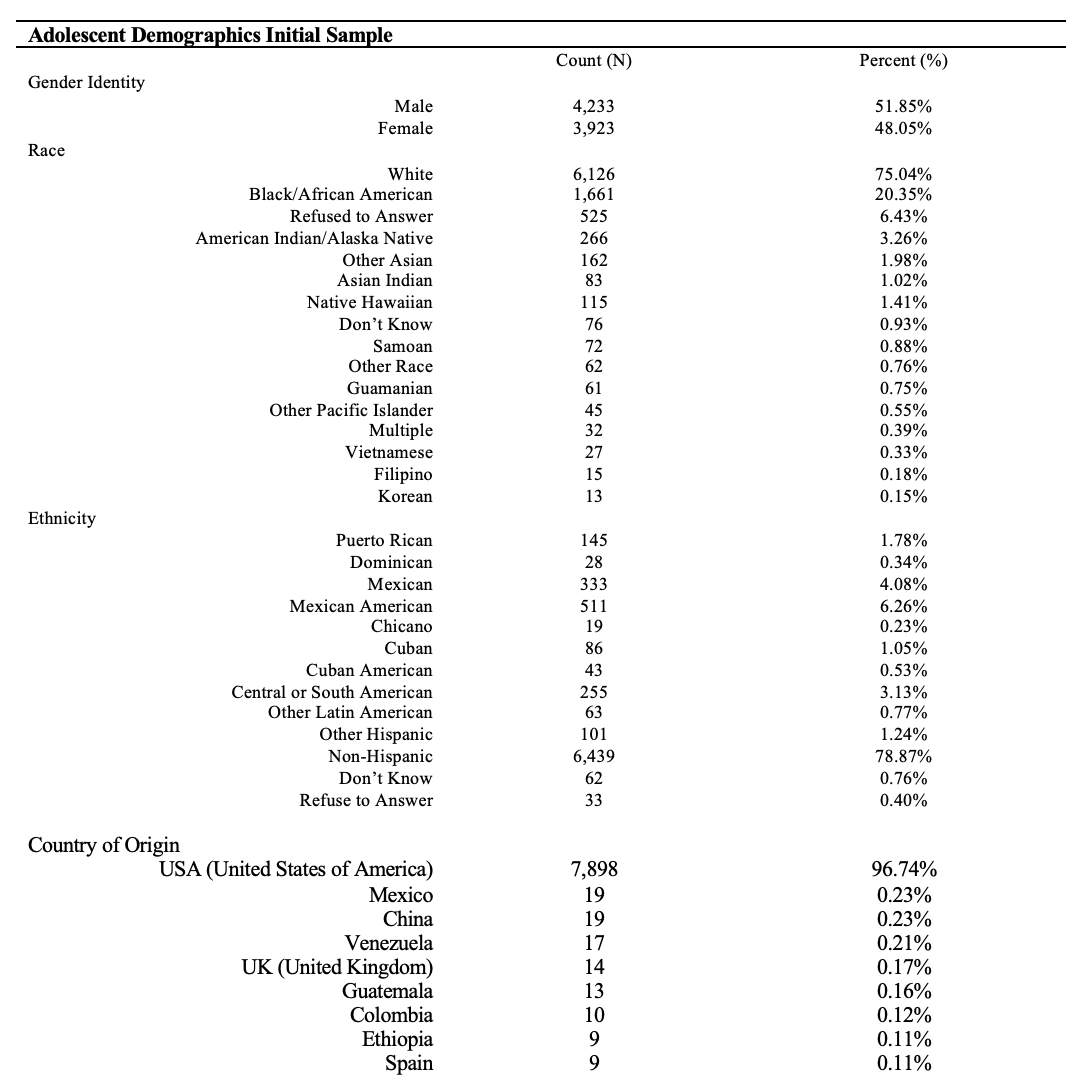

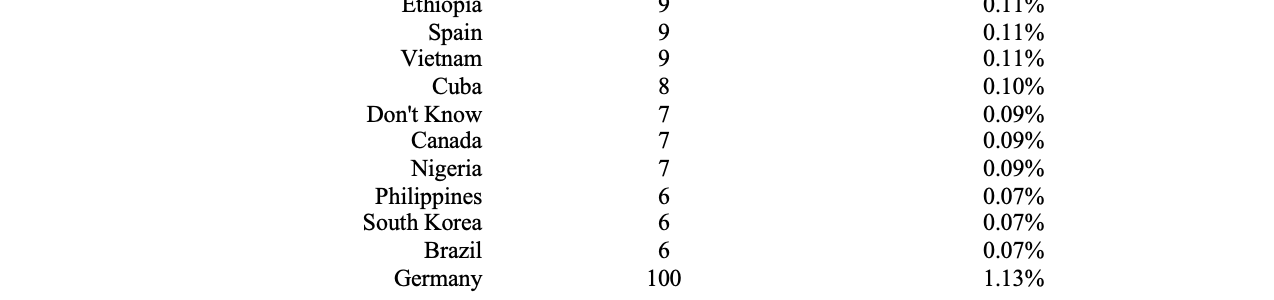

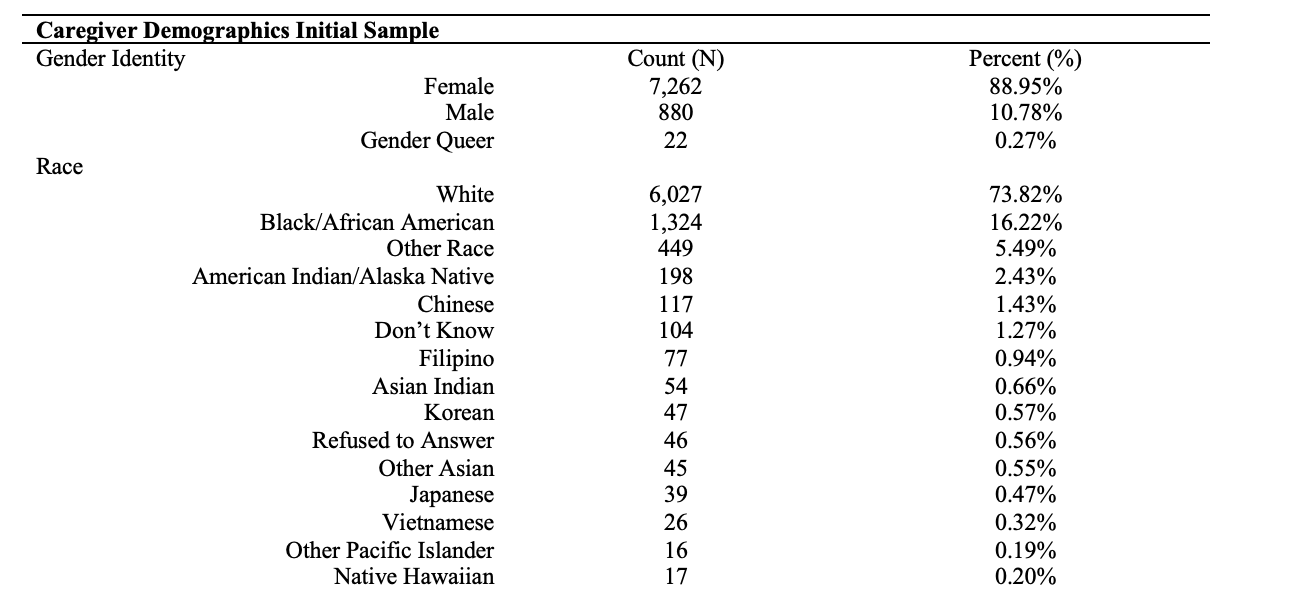

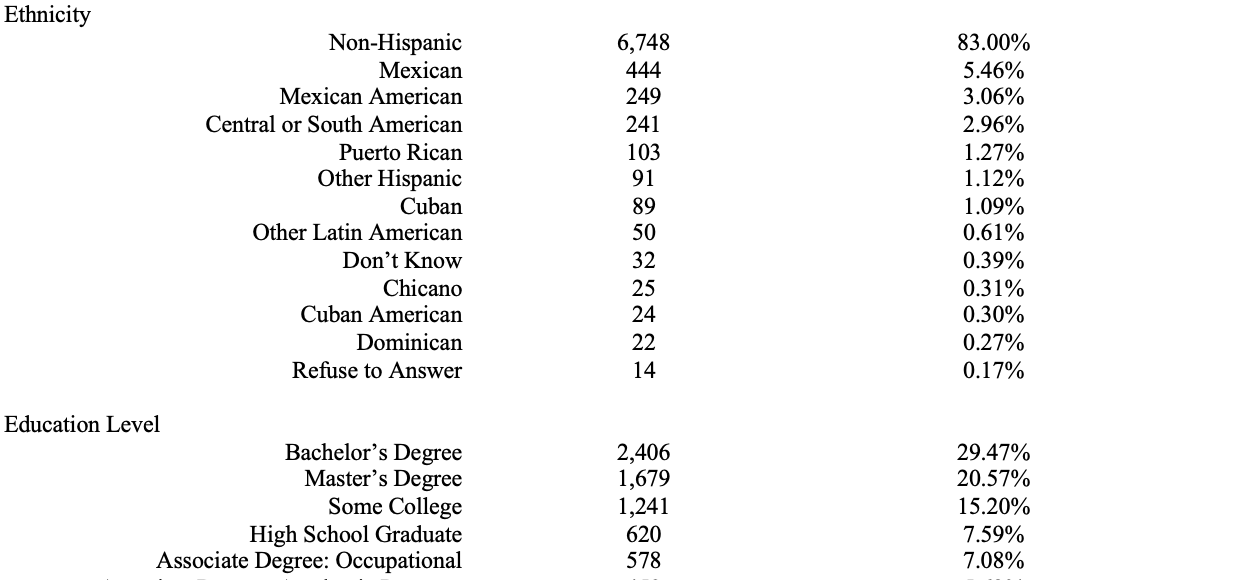

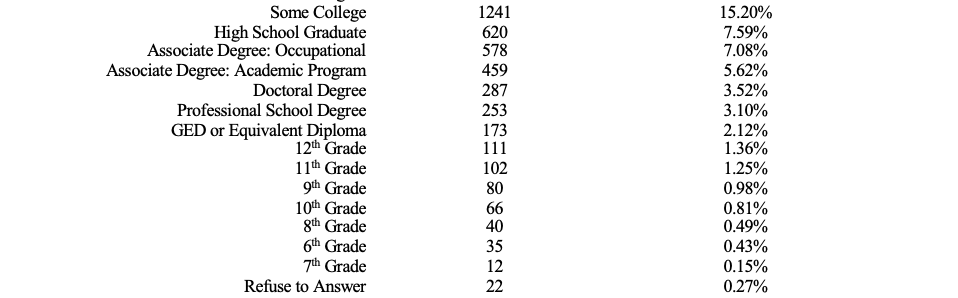

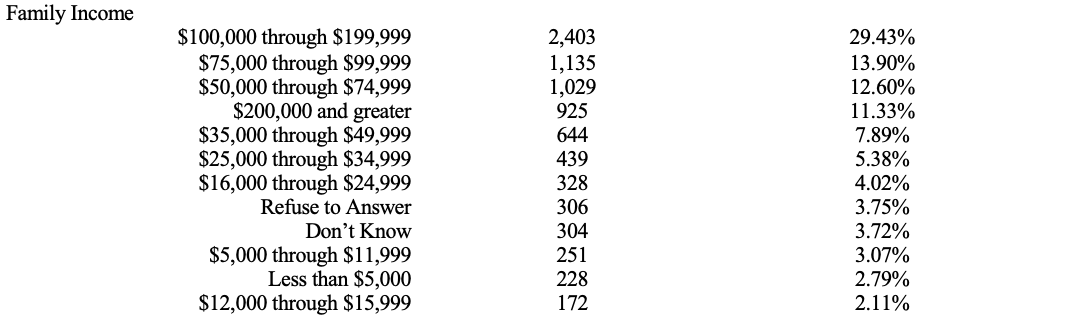

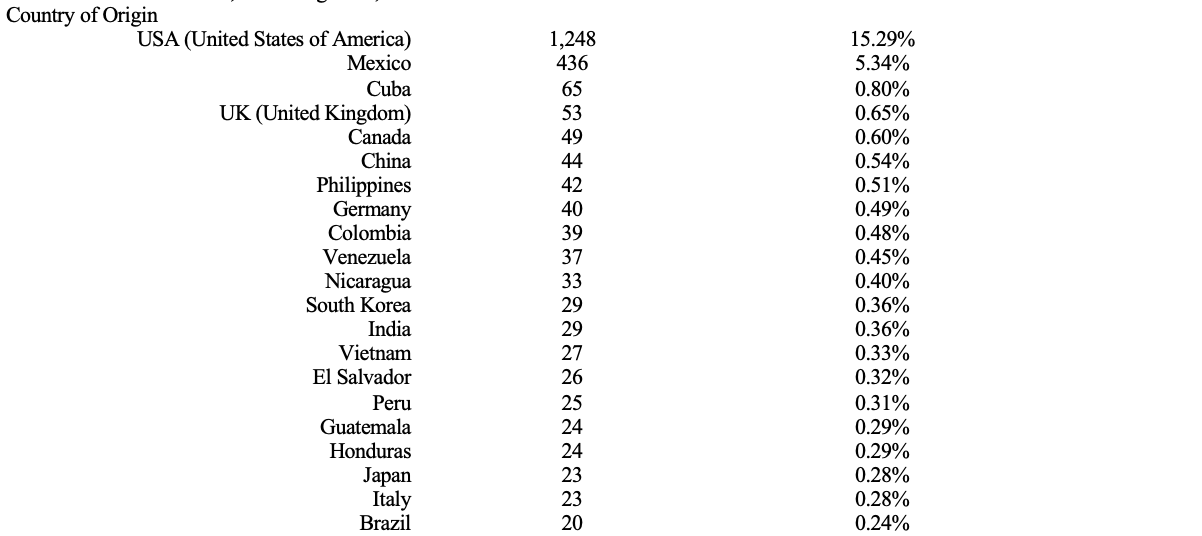

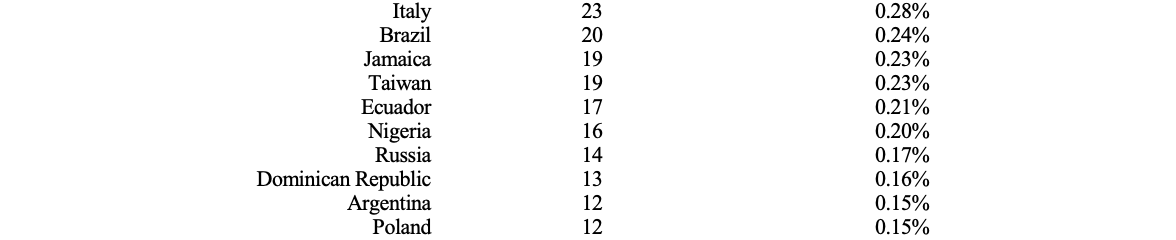

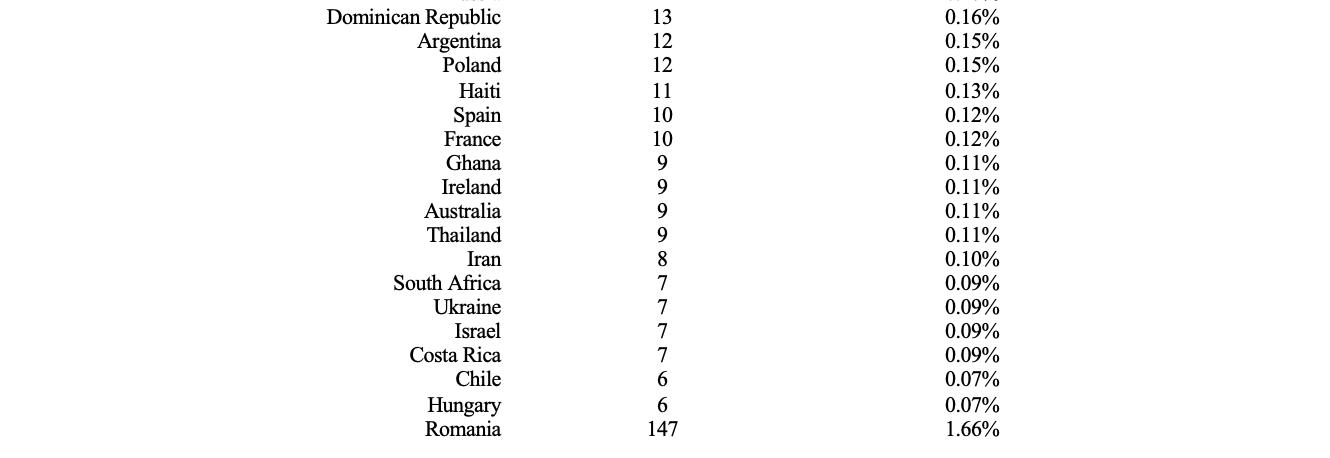

**Table S1. Full Demographics of MEIM-R Y3 Sample.** Variables which contained less than or equal to 5 participants were collapsed into reported categories to maintain anonymity (CPRD, 2024).

#### **Resting State Functional Connectivity**

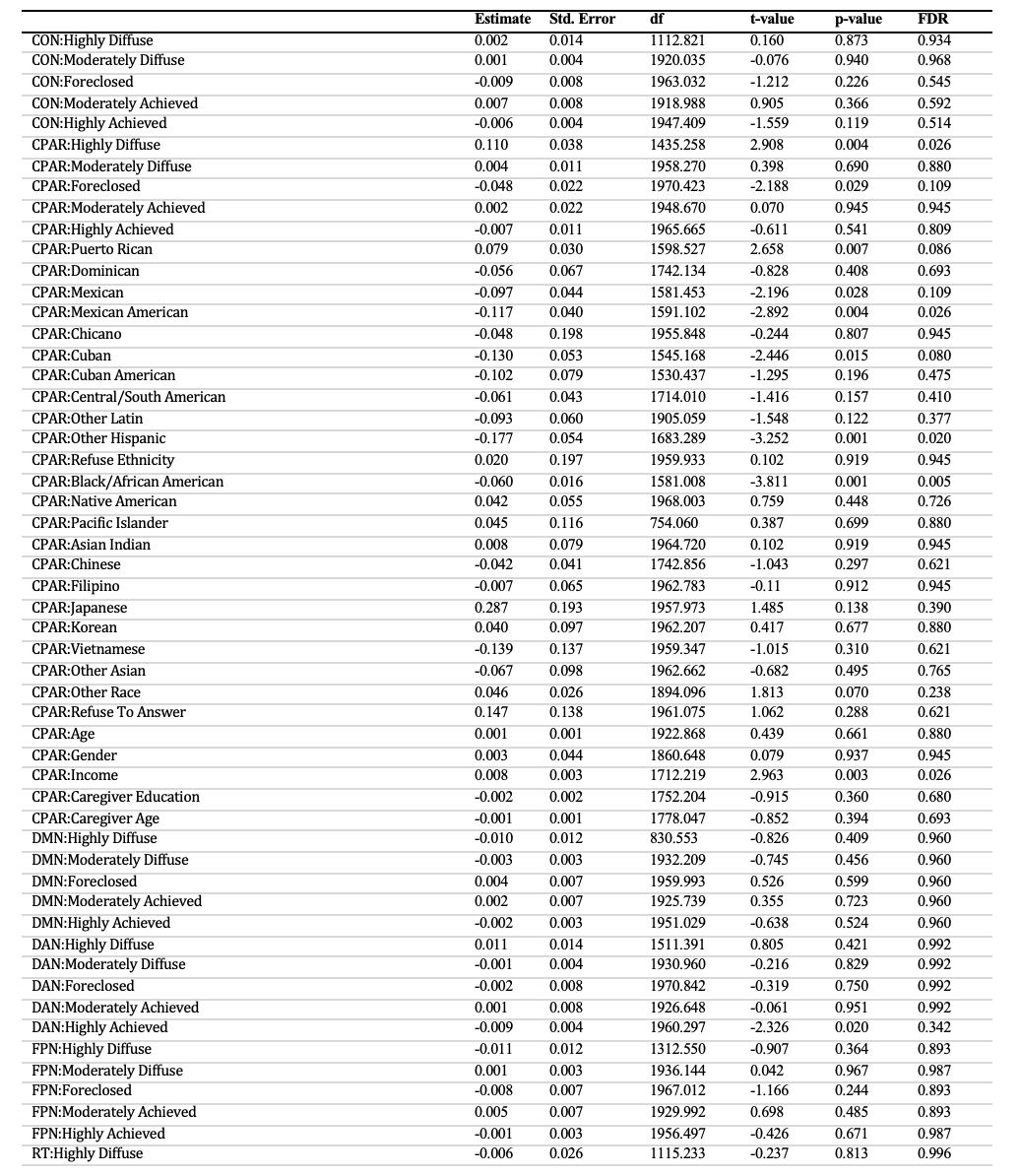

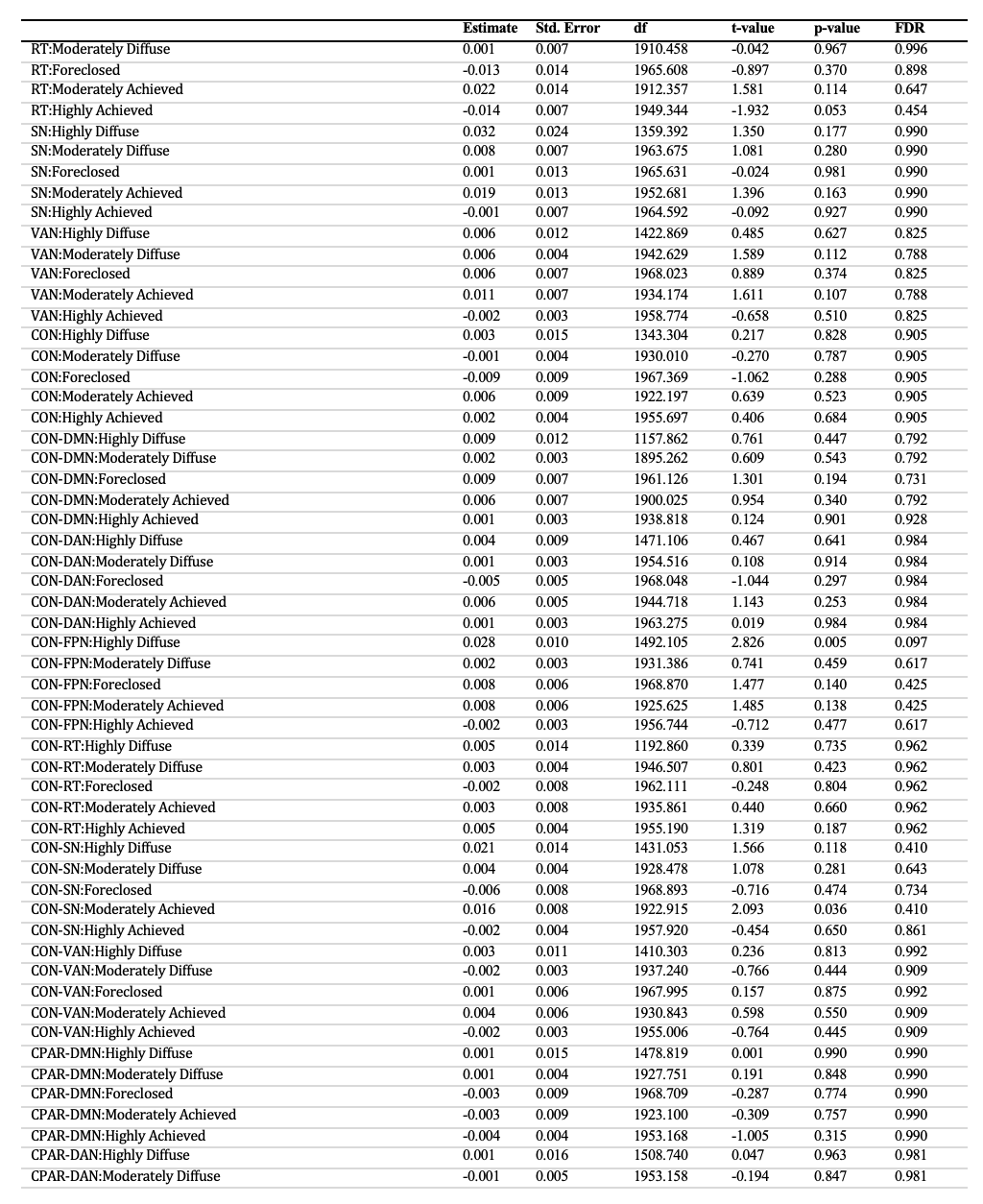

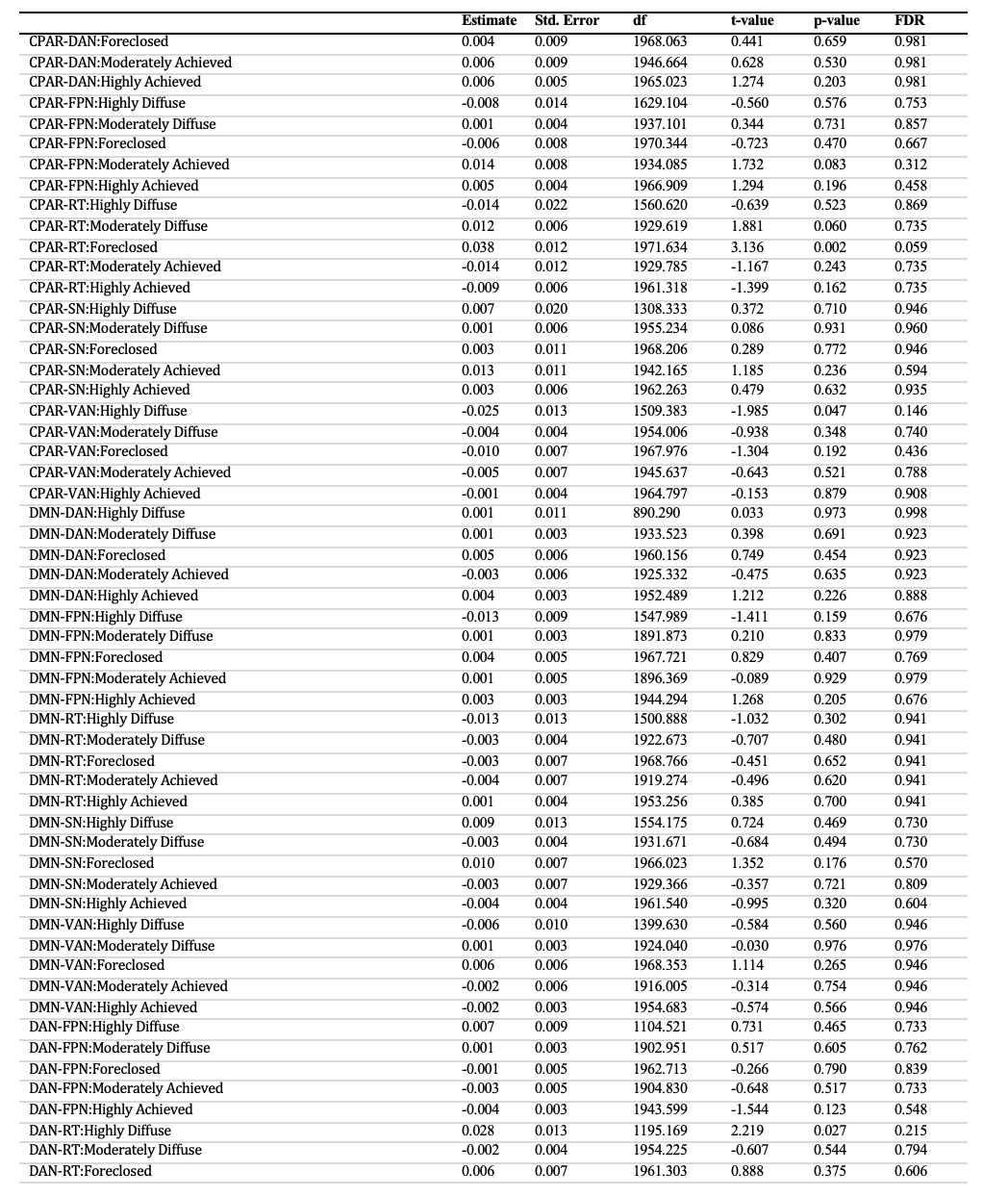

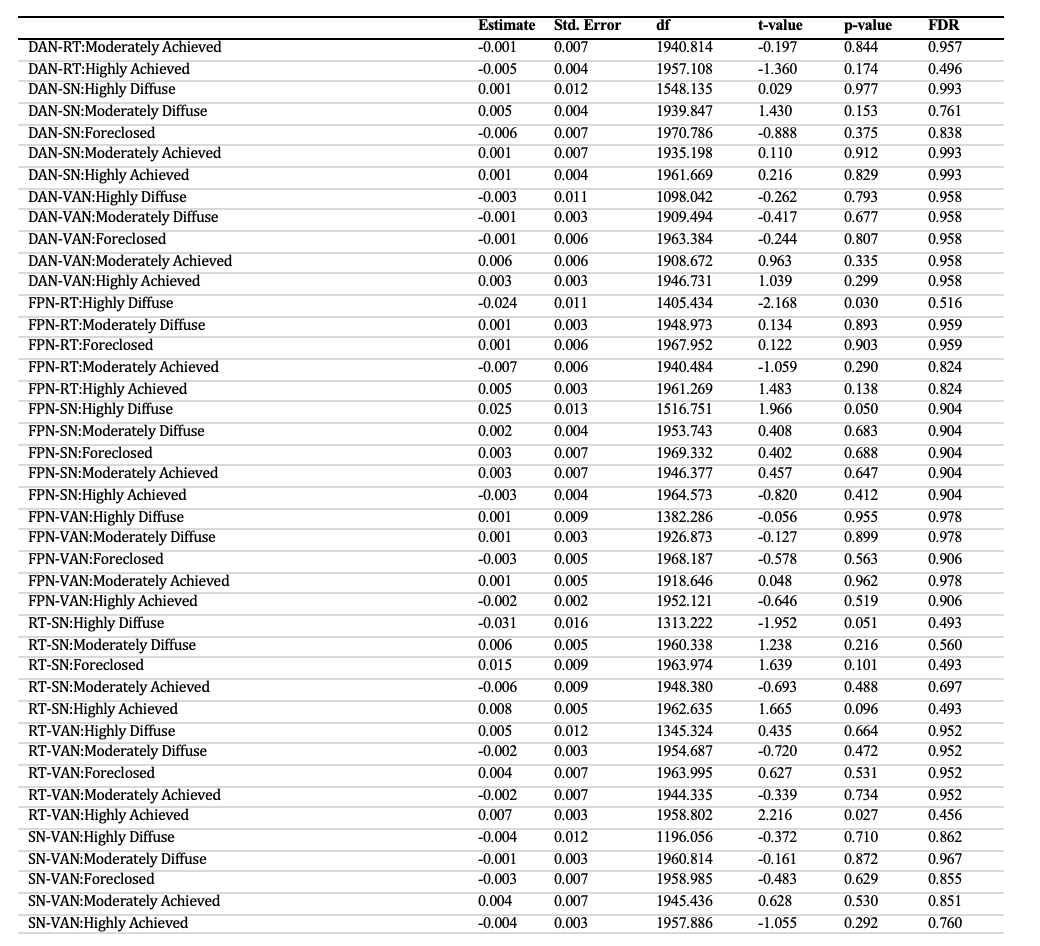

**Table S2. Resting State Functional Connectivity Results.** Statistical significance was determined by a p_FDR_–value of 0.05 or less. Degrees of freedom are denoted as df, standard error is denoted as Std. Error, and false discovery rate is denoted as FDR.

#### **Moderation of Perceived Discrimination**

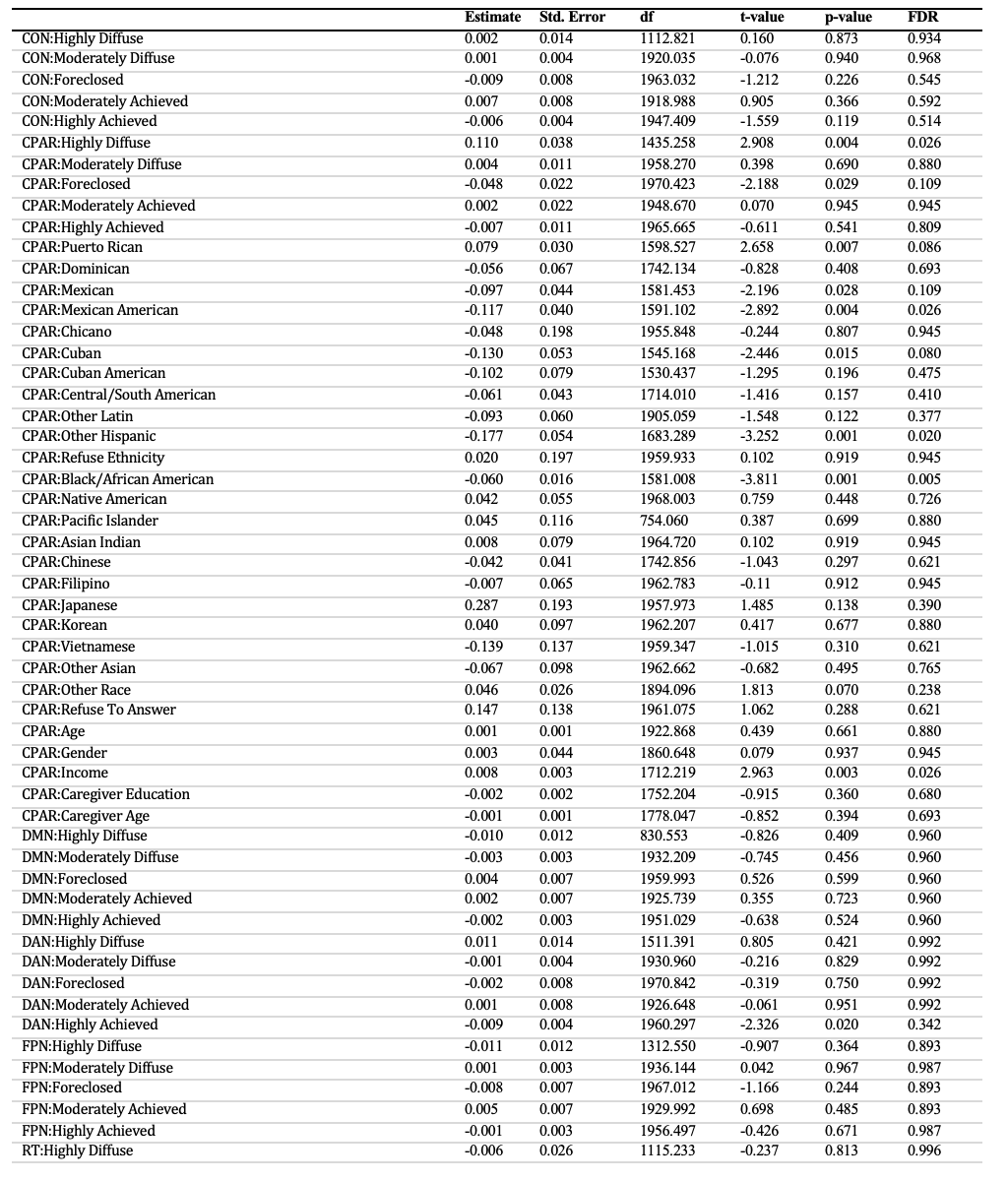

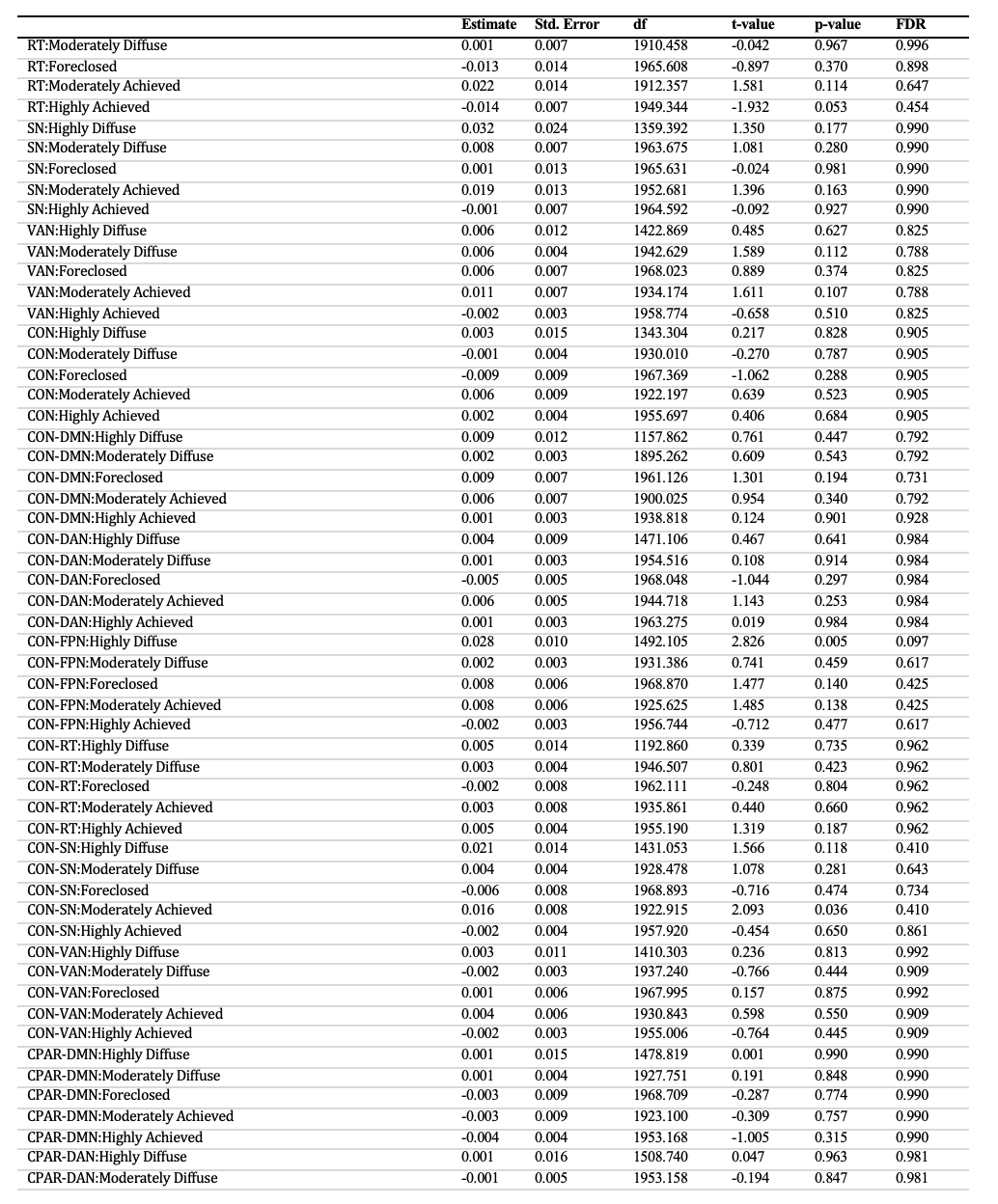

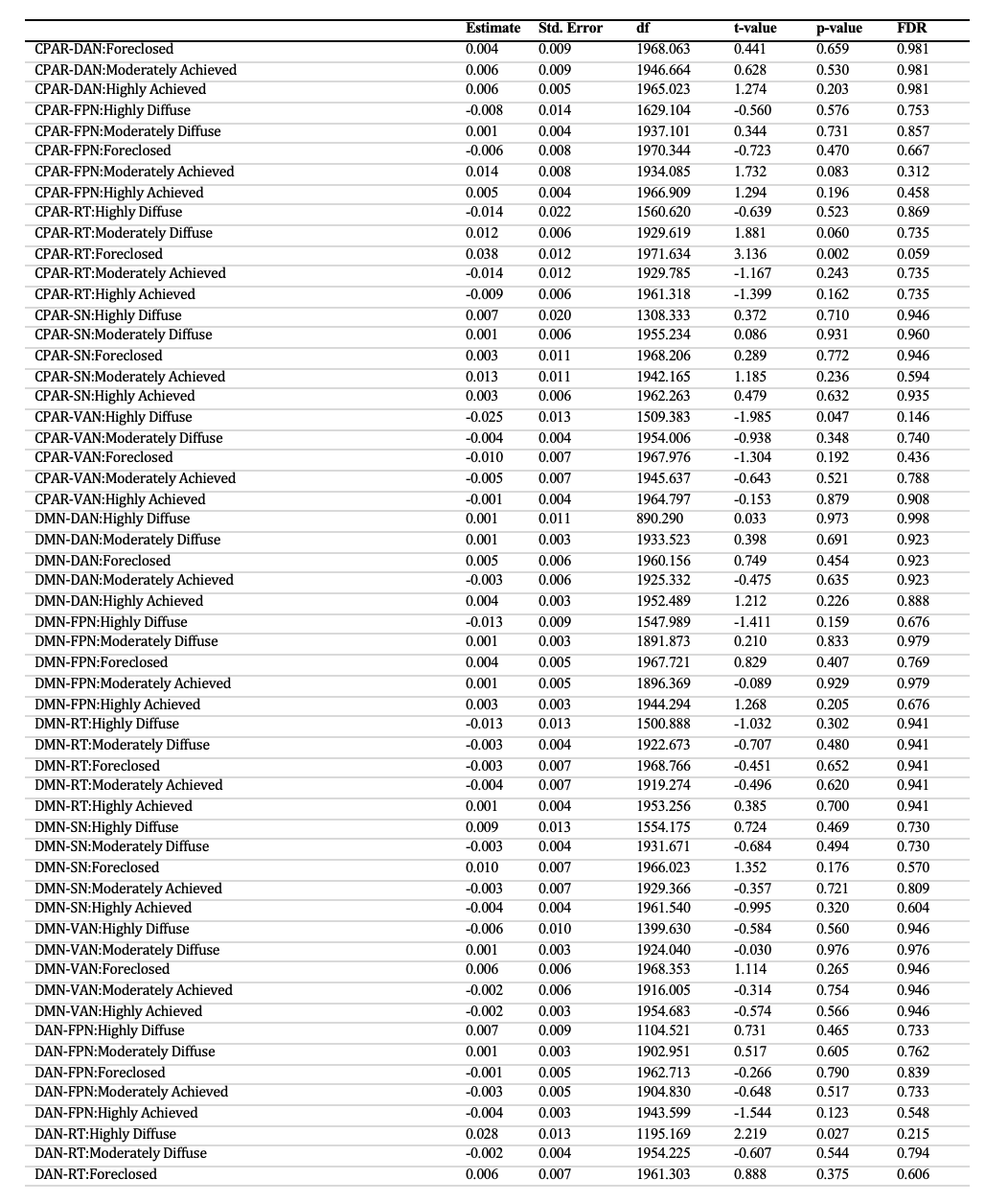

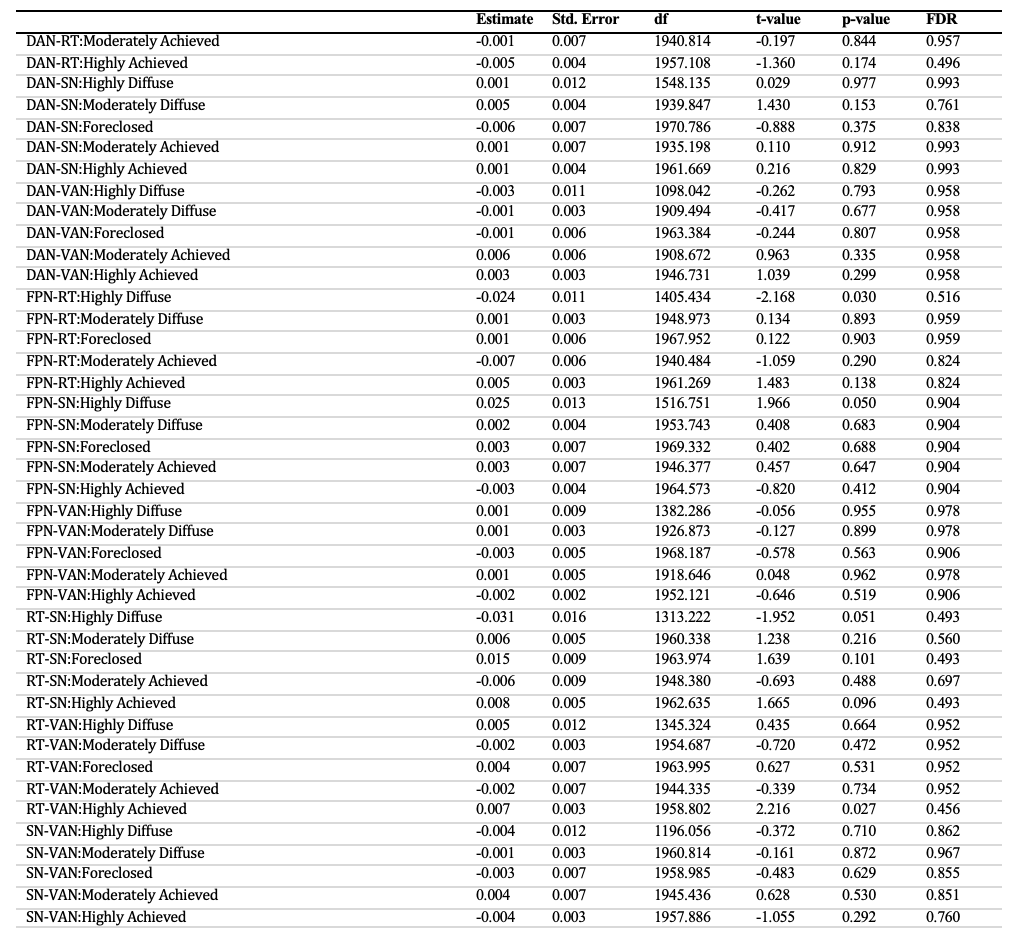

**Table S3. Moderation of Perceived Discrimination Results.** A moderation of perceived discrimination revealed within-CPAR significant ethnic identity profile associations. Statistical significance was determined by a p_FDR_–value < 0.05. Degrees of freedom are denoted as df, standard error is denoted as Std. Error, and false discovery rate is denoted as FDR.

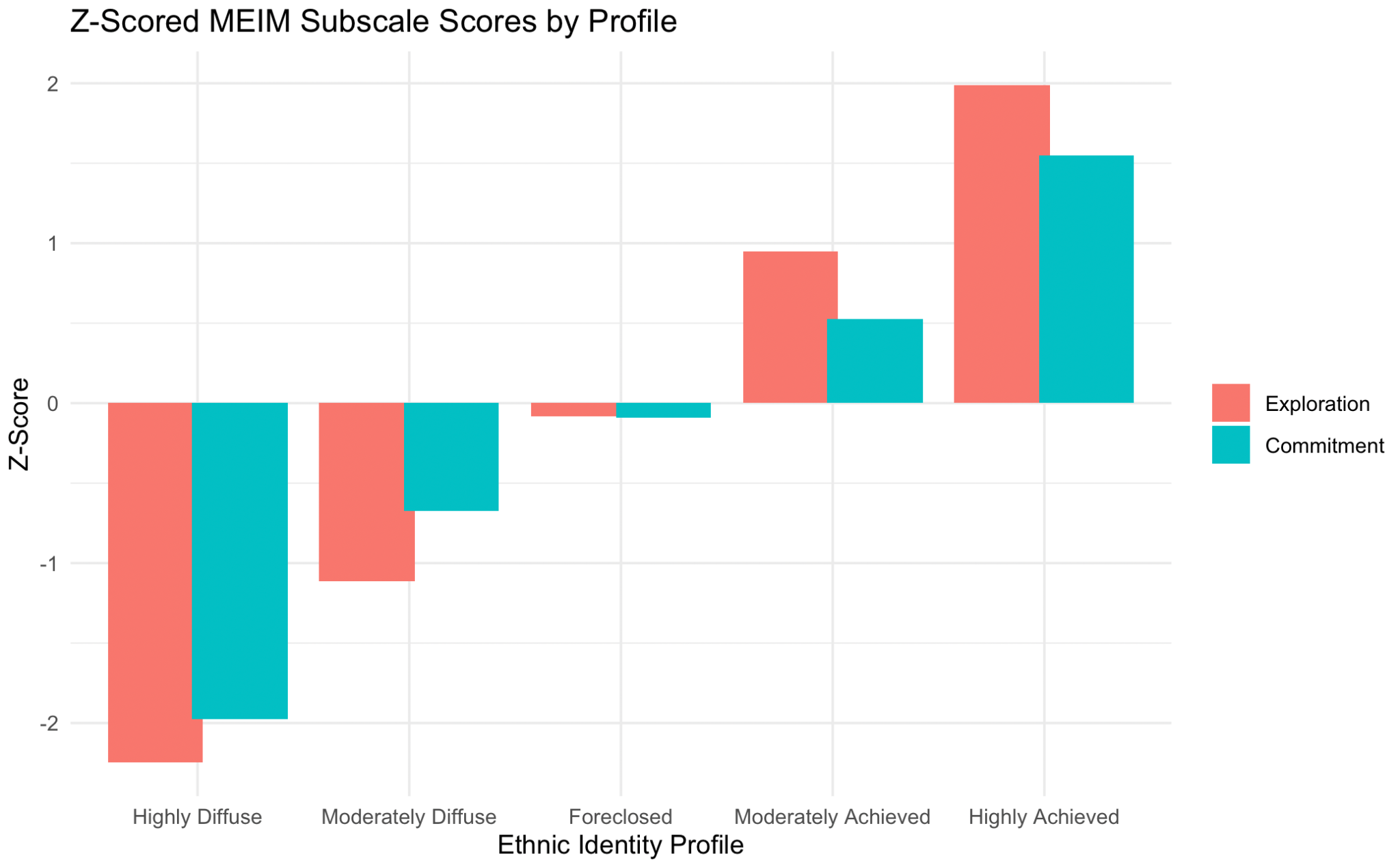

**Figure S1. Ethnic Identity Profiles by Z-Scores.**
